# Circuit mechanisms underlying chronic epilepsy in a mouse model of focal cortical malformation

**DOI:** 10.1101/2020.04.21.054148

**Authors:** Weiguo Yang, Anthony Williams, Qian-Quan Sun

## Abstract

**Highlights:**

- Ectopic interlaminar excitatory inputs from infragranular layers to layer 2/3 pyramidal neurons is a key component of the hyperexcitable circuitry
- Disrupted E/I balance was located far away from cortical malformations
- Dendritic inhibition from somatostatin interneurons play a key role in epileptogenesis
- Closed-loop optogenetic stimulation to activate remainder somatostatin interneurons irreversibly stops the spontaneous spike-wave discharges in vivo.

**In Brief:** Yang et al. report abnormal synaptic reorganization in an epileptogenesis zone in a mouse model of cortical malformation. The authors further demonstrate that spontaneous spike-wave discharges can be curbed by selectively activating somatostatin interneurons using close-loop fiber optogenetic stimulation to a small cortical region away from the microgyrus.

**Summary:** How aberrant neural circuits contribute to chronic epilepsy remains unclear. Using a mouse model of focal cortical malformation with spontaneous seizures, we dissected the circuit mechanisms underlying epileptogenesis. Spontaneous and optogenetically induced hyperexcitable bursts *in vivo* were present in a cortical region distal to (> 1mm) freeze-lesion induced microgyrus, instead of a region near it. ChR2-assisted circuit mapping revealed ectopic interlaminar excitatory inputs from infragranular layers to layer 2/3 pyramidal neurons as a key component of the hyperexcitable circuitry. This disrupted balance between excitation and inhibition was prominent in the cortical region distal to the microgyrus. Consistently, the synapses of both parvalbumin-positive interneurons (PV) and somatostatin-positive interneurons (SOM) to pyramidal neurons were maladaptive in a layer- and site-specific fashion. Finally, closed-loop optogenetic stimulation of SOM, but not PV, terminated spontaneous spike-wave discharges. Together, these results demonstrate highly site- and cell-type specific synaptic reorganization underlying chronic cortical epilepsy and provide insights into potential treatment strategies for this devastating neurological disorder.

## Introducion

A significant portion of intractable epilepsy in patients is intimately associated with the malformation of cortical development (MCD) (Guerrini et al., 1992; Taylor et al., 1971). Polymicrogyria (PMG) is the most common form of MCD in human epilepsy patients and often occurs as an isolated brain malformation (Leventer et al., 2008). A typical characteristic of human PMG patients is a 4-layered abnormal lamination surrounded by 6-layered cortical tissue (Dvorak and Feit, 1977; Dvorak et al., 1978). Surgical removal of microgyrus area usually generates good outcomes for only 50% of patients with intractable epilepsy due to PMG (Palmini et al., 1991). An effective treatment requires us to further understand how neural circuits are rewired into the aberrant network which leads to focal epileptogenesis (Guerrini et al., 1992), which remains largely unclear.

Dvorak et al. first induced experimental microgyrus in postnatal day zero rats via transcranial freeze lesion (FL) (Dvorak and Feit, 1977; Dvorak et al., 1978). Subsequent studies in brain slices found that epileptiform activities are capable of propagating over a long distance and were initiated in cortical regions adjacent to experimentally induced microgyrus, but not in the microgyrus itself (Jacobs et al., 1996; Luhmann and Raabe, 1996). However, electrographic or behavioral seizures were not found to be present in FL treated rats (Brill and Huguenard, 2010; Gibbs et al., 2008; Kellinghaus et al., 2007; Scantlebury et al., 2005), at least without a provoking event (Luhmann et al., 2014). We have recently developed a mouse FL model of MCD and demonstrated continuous spontaneous spike-wave seizures in both anesthetized and freely behaving mice (Williams and Sun, 2019; Williams et al., 2016). Noticeably, pathological high frequency oscillations, which have been used as a reliable biomarker for identifying seizure-generating zones (Bragin et al., 2004; Worrell et al., 2004) in resection surgical treatment of refractory epilepsies (Haegelen et al., 2013), were elevated in the superficial cortical area distal to the microgyrus in the FL mouse model (Williams and Sun, 2019). These studies allowed us to interrogate the circuit mechanisms underlying chronic epileptogenesis in epileptic mice.

To tackle this question, we used ChR2-assisted circuit mapping (CRACM) to identify aberrant synapses and to assess their E/I balance (Yang and Sun, 2018). We then combined optogenetic stimulations with linear microelectrode array recordings *in vivo* and found that the epileptogenesis site was remote to the microgyrus, which was confirmed by CRACM analysis in cortical regions proximal or distal to microgyrus. Strikingly, an unexpected aberrant L5B → L2/3 input, located distal from the microgyrus, contributed significantly to the maladaptive hyperexcitable circuit. Concurrently, two major cortical interneurons (INs), PV and SOM, experienced site-specific reorganization that was consistent with their roles in the E/I imbalance. Closed-loop optogenetic stimulation of SOM, but not PV, terminated spontaneous spike-wave discharges. Together, these experiments indicate that balanced synaptic excitation and inhibition is interrupted in a site-, layer- and cell type-specific manner in FL-induced epileptic mice.

## Results

### Reversed layer-specific E/I balance in epileptic mice, primarily due to *de novo* inter-laminal excitatory inputs to L2/3 neurons

To uncover the cellular mechanisms in epileptic circuits, we sought to examine canonical circuits within the barrel field of the somatosensory cortex (vS1), but not in the microgyrus itself, using CRACM approach in brain slices (Mao et al., 2011; Petreanu et al., 2007; Petreanu et al., 2009; Yang and Sun, 2018). We used CRACM to map the strength and spatial pattern of intracortical synaptic inputs to both L2/3 and L5B pyramidal cells (PCs), which are the major cortical outputs to other brain regions (Aronoff and Petersen, 2006; Harris and Shepherd, 2015; Li et al., 2015). Injected AAV-ChR2 virus transfected both excitatory and inhibitory neurons in vS1, typically covering 2-4 barrel columns (Figures 1A and 1C, left, Methods). This allows us to study the effects of optogenetic activations of local synaptic inputs to the recorded PCs. Stimuli were delivered on a photo-stimulation grid (16 x 12, 75 µm spacing) covering a 1.1 mm^2^ area surrounding the recorded neuron (Figures 1A and 1B).

**Figure 1.**
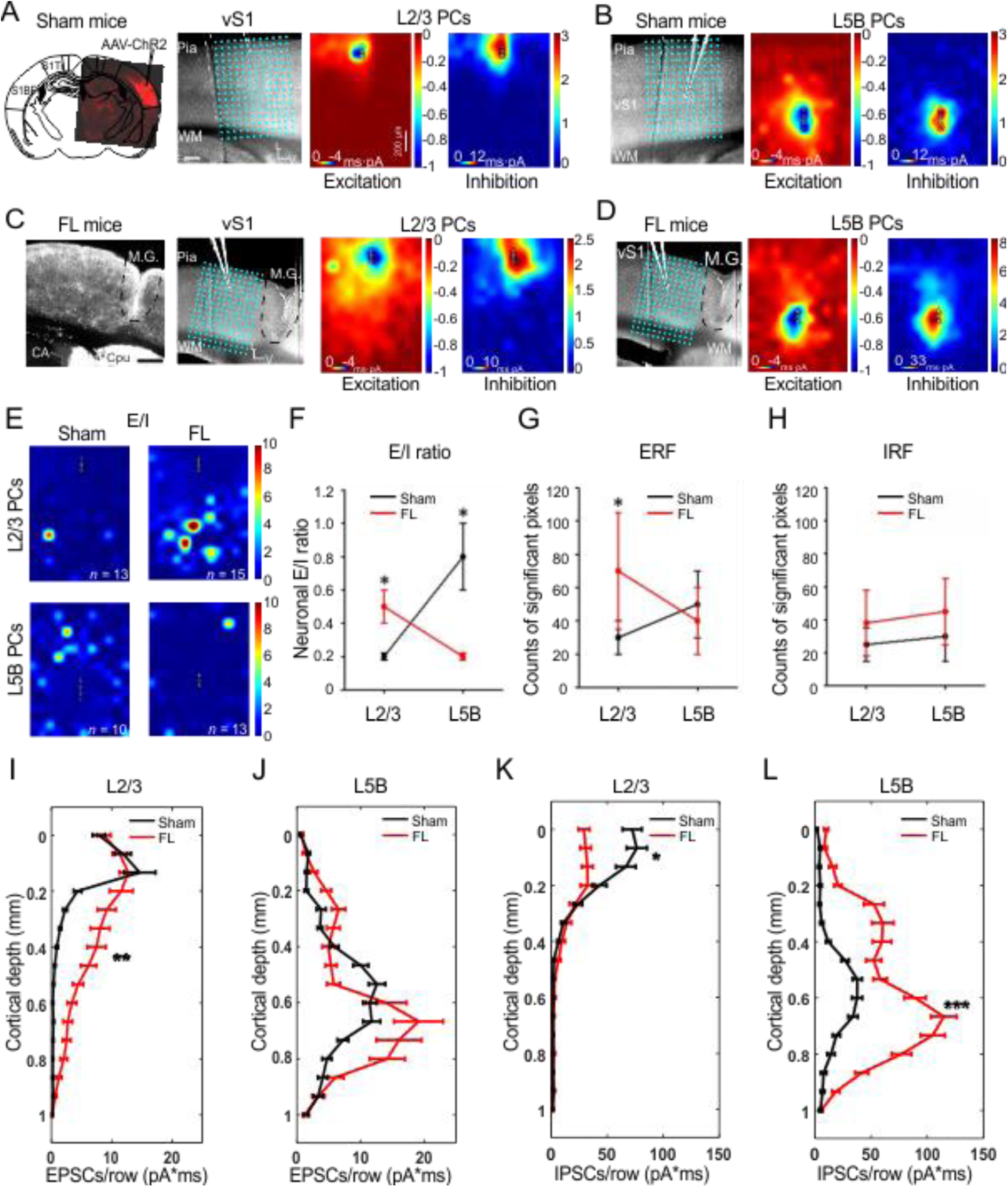
Aberrant synaptic inputs to superficial PCs in S1. **(A-D)** CRACM of EPSCs and IPSCs in S1 L2/3 (**A,C**) and L5B (**B,D**) PCs in sham mice (**A,B**) vs. FL mice (**C,D**). In each condition (**A-D**), an overlay of a brain slice with a 16-by-12 photo-stimulation grid illustrates the photo-stimulation and recording sites for each group. Left (**A,C**): two confocal images of AAV-ChR2 virus expression pattern in vS1 in sham and FL, respectively. Right (**A-D**): CRACM maps of excitatory synaptic inputs (“Excitation”) and inhibitory synaptic inputs (“Inhibition”) in L2/3 PCs (neuron and mice numbers see **E**). Triangles indicate neuronal soma locations in this and following panels and figures. Note the excessive excitatory input to L2/3 PCs in FL mice (compare Excitation maps in a and c). Scale bar, 200 µm. **(E)** E/I heat maps obtained by calculating EPSC/IPSCS ratios in each of the 192 sites (16 X 12 pixels) in L2/3 (n=13 neurons for sham and n=15 for FL) and L5B PCs (n = 10 neurons for sham and n=13 for FL) in sham (n=5 mice) vs. FL mice (n=5 mice). **(F)** E/I ratios were obtained by calculating the ratios of the summed EPSCs charges over summed IPSCs charges in heat maps. The E/I ratio in L2/3 PCs in FL group was significantly larger than that in sham group (unpaired t-test, P=0.027); meanwhile, the E/I ratio in FL L5B PCs was significantly reduced compared to that in sham counterparts (unpaired t-test, P=0.012). **(G)** Excitatory receptive field (ERF), obtained by counting significant EPSC pixels (>2 x sd) in each group. The ERF in sham L2/3 PCs was significantly higher in FL group than their counterparts in sham group (unpaired t-test, P=0.037); the ERF in L5B PCs was similar across groups (unpaired t-test, P=0.83). **(H)** Inhibitory receptive field (IRF), obtained by counting significant IPSC pixels in each group. No significant IRF change was found in both L2/3 and L5B PCs across groups (unpaired t-test, P=0.71 for L2/3 and P=0.63 for L5B). **(I-L)** Laminar distribution of average charges or strength of EPSCs (**I,J**) and IPSCs (**K,L**) for L2/3 and L5B PCs from sham (black) and FL (red) mice. For L2/3 PCs, the EPSCs charge was significantly increased in FL group (unpaired t-test, P=0.0013). In contrast, L5B EPSCs charge in FL group was similar to that in sham group (unpaired t-test, P=0.134). However, for IPSCs charge, the layer specific difference was roughly reverse between FL and sham groups (L5B, unpaired t-test, P=0.000; L2/3, P=0.044).

Polysynaptic input to recorded neurons typically maximized at the perisomatic region (Figure 1, Figure S1). Excitatory or inhibitory postsynaptic potentials (PSP) displayed shorter latency around perisomatic regions (Figure S1D). Application of TTX and 4-AP would eliminate synaptic input from intermediate neurons and reveal somacentric and monosynaptic input (Figures S1E and S1F), consistent with previous results (Petreanu et al., 2009). We used an excitatory or inhibitory “receptive field” (ERF/IRF), which consisted of pixels with significant synaptic events (see Methods), to visualize the spatial patterns of PSC events or their changes. TTX eliminated ~70% of ERF of L2/3 PCs (P<0.01, n = 5) and ~60% of ERF of L5B PCs (P<0.05, n = 5).

In the L2/3 PCs of sham mice, optogenetically evoked synaptic responses were mainly restricted to supragranular layers (Figure 1A, Excitation and Inhibition maps). The median E/I ratio across pixels (pixel E/I ratio) was 0.19 (*n* = 13 neurons, Figure S2A). Different stimuli intensities led to similar neuronal E/I ratios (summed E and I across pixels, see Methods, Figure S2), suggesting that the E/I ratios are intrinsically regulated and balanced, as previously reported in a normal mouse cortex (Wehr and Zador, 2003; Xue et al., 2014; Yang and Sun, 2018). Simultaneously stimulating the majority, if not all, of the synaptic inputs to the recorded neurons using whole-field LED illumination, showed a similar result (Figure S2F). Importantly, extensive supragranular and ‘ectopic’ infragranular excitatory inputs to L2/3 PCs were revealed (compare the Excitation maps in Figures 1A vs 1C). As a result, numerous ‘hot pixels’ (meaning higher E/I ratios) appeared within the E/I ratio map (Figure 1E, top right), mainly in the infragranular layers, which was in sharp contrast to the sham mice (Figure 1E, top left). The neuronal E/I ratios were significantly elevated in L2/3 FL group (Figure 1F, P<0.05); the ERF showed larger ERF in L2/3 FL group (Figure 1G, P<0.05). Consistent with above findings, the layer-origin of the significant increases in optogenetically evoked EPSPs to L2/3 PCs spans from lower L2/3 all the way to L6 (Figure 1I, P<0.01). Also, the average median value of pixel E/I ratios of L2/3 PCs showed dramatically significant increase from 0.19 in sham to 0.63 in FL (Figure S2C, P=0.0026), suggesting that the synaptic E/I balance in FL treated L2/3 PCs had been maladaptively disturbed in the following manner: higher excitatory inputs from both inter-laminar and inter-columnar S1 regions, particularly *de novo* excitatory inputs from infragranular layers (Figure 1j).

In sham treated mice, L5B PCs were characterized with stronger excitatory inputs and higher E/I ratio compared with that of L2/3 PCs from the same columns (median 0.41 vs. 0.19, Figure S2). Like L2/3 PCs, L5B E/I ratios maintained a similar value under two different intensities (Figure S2B). Surprisingly, FL L5B PCs received stronger inhibition compared to their sham-counterparts (Figures 1B and 1D, Inhibition maps). Despite slight increases in IRF (Figure 1H), the IPSC strength at L5B PCs was much stronger (Figure 1l), leading to significantly lower E/I ratios in FL group (Figure1E, Figure S2, median 0.13 vs. 0.41). In addition, neuronal E/I ratios decreased with higher laser intensity in this group (0.08, *P* = 0.031, *n* = 9 neurons, Figure S2F), suggesting a hyper-inhibitory innervation of L5B PCs in FL mice. Thus, the E/I ratios in FL L2/3 PCs were several folds higher than their sham-counterparts, whereas the E/I ratios in FL L5B PCs were significantly lower than their sham-counterparts (Figure 1F, Figures S2A 2E).These results indicate a drastic rewiring has occurred in the cortical circuits of mice with chronic epilepsy associated with MCD.

### The hyperexcitable site was located in supragranular layer distal from center of microgyrus

Previous *in vitro* studies using the FL rat model found hyperexcitability adjacent to the FL-induced microgyrus (Jacobs et al., 1999). However, it is unclear how cortical circuits are maladaptively reorganized in mouse MCD models and the relationship to the hyperexcitable site *in vivo*. Using *in vivo* microelectrode array recordings, we found that not only were extracellular waveforms altered in FL animals, there was a spatial change in both local field potential (LFP) and multi-unit activities (MUA) relative to the distance from the microgyrus (Figure 2A, Figure S3C). As such, FL data was further divided into regions either ‘distal (>1 mm)’ or ‘proximal (<1 mm)’ to the center of microgyrus as determined from histological analysis of post-mortem brain tissues (Figure S3B).

**Figure 2.**
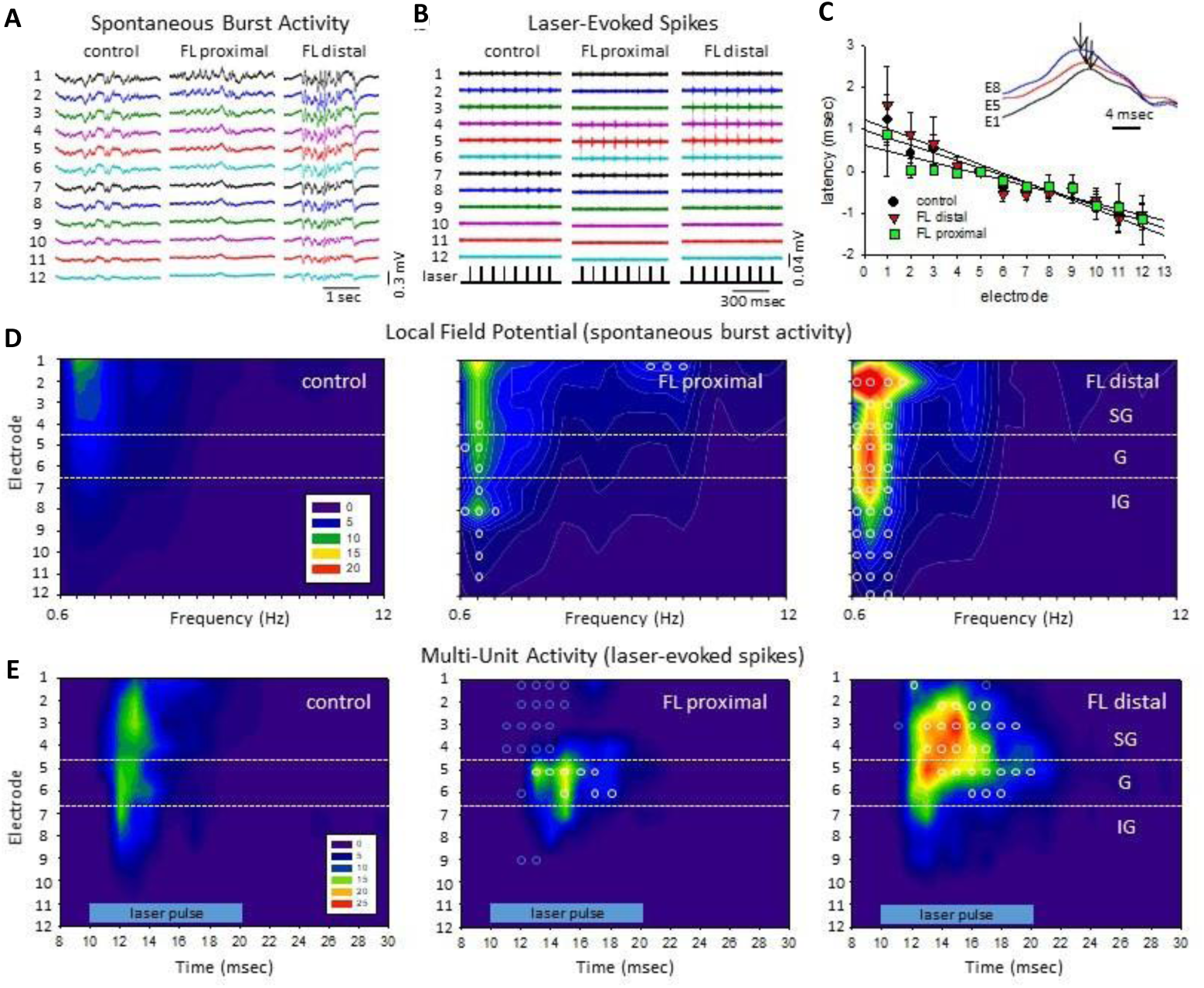
Site-specific hyperexcitability revealed by measuring spectral power of spontaneous burst activity and laser-evoked MUA in epileptic mice. **(A)** Exemplar unfiltered LFP recordings showed much stronger burst activities in distal region (>1 mm from the FL microgyrus) from S1BF of FL mice compared to those in anesthetized sham-treated mice or proximal (<1 mm from the FL microgyrus) region of FL mice. **(B)** Exemplar laser-evoked MUA from control, FL-proximal and FL-distal groups. **(C)** Shift in latency of burst waveforms computed from the peak offset of the cross-correlation values between electrodes (see **Methods**, r2 = 0.913 (control), 0.874 (FL-distal), and 0.879 (FL-proximal)), inset shows the corresponding shift in waveform peaks across 3 electrodes (E1, E5, and E8, black arrows indicate peak deflection). Importantly, note that the shortest latency is in the infragranular layers (E8) suggesting the origin of, at least partial, synaptic input to superficial layers. **(D)** Composite heat maps of power spectral density (PSD) values from spontaneous burst activities in the three groups. **(E)** Composite heat maps of laser-evoked MUA counts across cortical lamina (horizontal blue bar, laser intensity level 8). **D-E**, Heat maps were constructed from the average values of spike counts from each experimental group; control (n=6 mice), FL-distal (n=4 mice) and FL-proximal (n=4 mice). Horizontal white dotted lines indicating the transitions between supragranular (SG), granular (G), and infragranular (IG) layers. Blue indicates lower spike counts while red indicates higher counts. Open circles indicate significant differences compared to control groups (P<0.05, Holm-Sidak post-hoc analysis).

To determine the precise loci of hyperexcitable waves *in vivo*, we used two complementary approaches: anesthesia and optogenetically evoked bursts discharges (see Methods). First, we measured the latency of optogenetically evoked bursts *in vivo*, as this can be accurately measured with regard to the time of laser stimulations (Methods). Under isoflurane anesthesia, extracellular waveforms of laser-evoked burst activity typically progressed from deeper to more superficial layers of the cortex, (Figure 2c) as indicated by the temporal shift in waveform peaks of the recorded LFP signals across layers (Figure 2C, inset). An average delay of 2-3 ms was observed in the supragranular layers compared to the infragranular layer (Figure 2C). The rapid progression of the burst waveform from the deeper to more superficial cortical layers was also associated with a significant increase in total signal power (0-20 Hz) in the supragranular cortical layers (P<0.05, ANOVA) only for the FL groups (P<0.05, Holm-Sidak post-hoc analysis). Next, we examined anesthesia-induced spontaneous bursts discharges, which we have found previously to be significantly enhanced by FL (Figure 2A)(Williams et al., 2016). Further analysis indicated significant differences in the spectral power and the frequency spectrum of these abnormal burst-discharges associated with the FL-treatment (P<0.05, multivariate ANOVA) as displayed in the laminar heat maps of Figure 2D. In comparison, the strongest increases in spectral power were observed in the supragranular layers of the FL-distal group, particularly in the delta frequency range (0-4 Hz). Last, because evoked MUA is the gold standard for accurate mapping of the origin of evoked bursts/waves, we examined laminar and lateral location of optogenetically induced bursts and found that optogenetic-evoked MUA showed significant increases in spike count and lasted much longer in the upper cortical layers of FL-distal group (Figure 2E, P<0.05, multivariate ANOVA). Interestingly, a significant reduction in spike count was also observed across some FL-proximal regions, particularly the supragranular layers (Figure 2E). Taken together, this *in vivo* data indicates that the distal region to the microgyrus is a likely locus for epileptogenesis and that epileptogenesis may originate from the infragranular layer.

### Identifying the location and sources of the hyperexcitable foci within the malformed cortex

Next, we asked how the previously mentioned hyperexcitability aligns with cortical regions proximal or distal to microgyrus at the cellular and synaptic level. We further divided the FL data in the Figure 1 into two groups: FL-proximal and FL-distal (Figures 3A-D, Figure S4). Importantly, L2/3 PCs in the distal group displayed a significantly larger E/I ratio than both the proximal group (Figure S4c, E/I ratio = 1.1 vs 0.3, *P*<0.01) and the sham group (Figure 3E, E/I ratio = 1.1 vs 0.2, *P*<0.01), respectively. Comparisons of the neuronal E/I ratios between proximal vs. distal groups under two laser intensities supported the findings mentioned earlier (Figure S4E). The ERF size in L2/3 PCs increased significantly compared to both the distal and sham groups (Figure 3F, *P*<0.01). The ERF and IRF sizes for L2/3 PCs were comparable in sham and proximal groups; however, ERF size outnumbered IRF size in distal group (P<0.01, Fig 3F, left panel). Consistently, the EPSC strength at L2/3 PCs was also significantly increased (Figure 3G, P<0.05); meanwhile the IPSC strength was reduced sharply when compared to sham or distal groups (Figure 3H, P<0.05). Importantly, the source of the ERF to L2/3 PCs were infragranular layers (Figure 3G, left panel). These results indicate that the epileptogenesis foci were mainly located at cortical regions distal (>1 mm) to center of the microgyrus, while the L2/3 PCs in the proximal (0-1mm) microgyral area received balanced excitation and inhibition compared to distal group.

**Figure 3.**
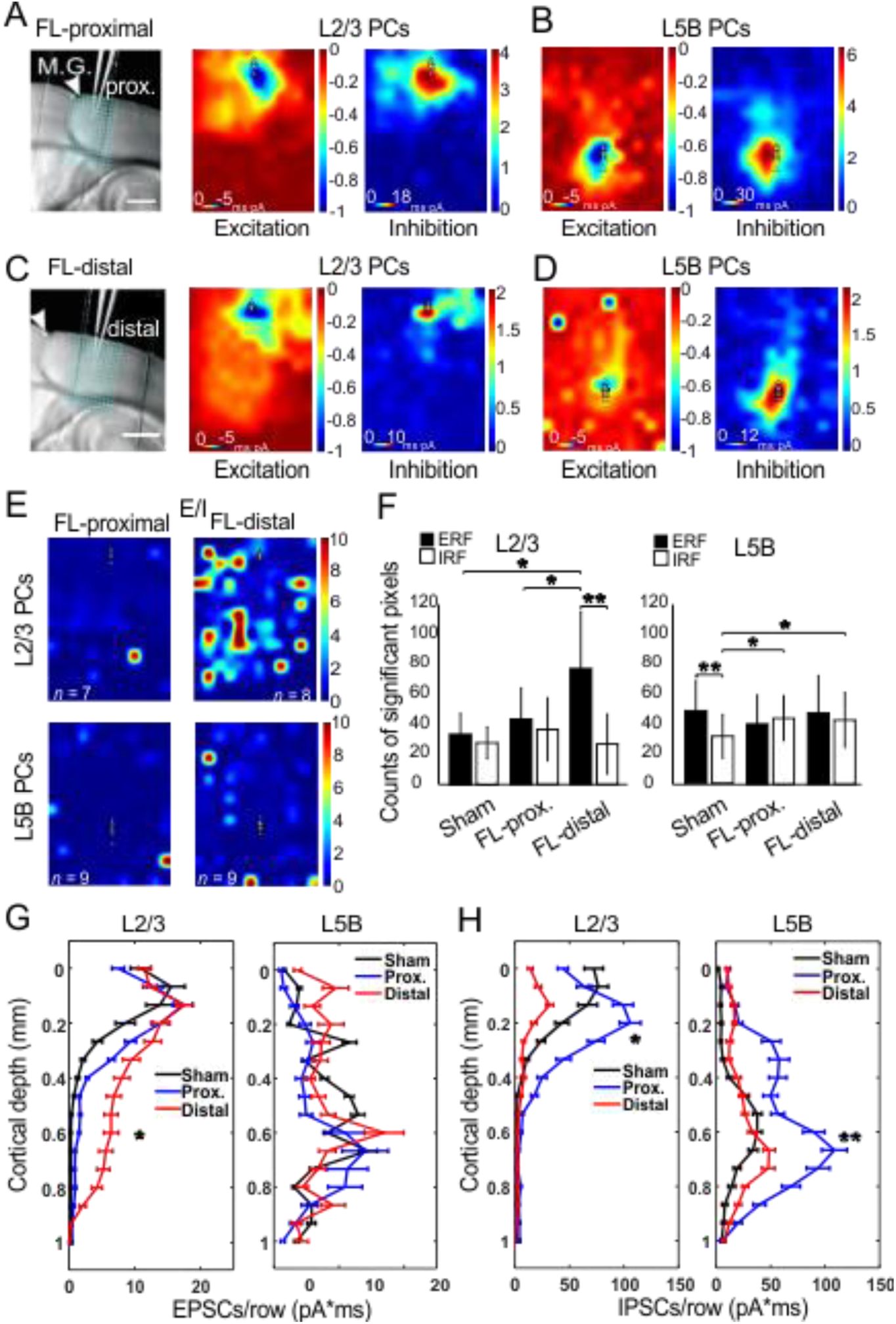
Site- and layer-specific synaptic inputs measured by CRACM. **(A-B)** CRACM of local vS1 excitatory and inhibitory synaptic inputs to L2/3 or L5B PCs in a region which were proximal to the M.G. **(C-D)** Same as above but for the FL-distal region. The excitatory input to distal L2/3 PCs was more extensive compared to its proximal counterpart, particularly in infragranular layers (compare Excitation maps in **A,C**); meanwhile the inhibitory input to distal L2/3 PCs was much smaller compared to its proximal counterpart (compare Inhibition maps in **A,C**). **(E)** E/I heat maps in L2/3 (n=7 neurons for FL-proximal and n=9 for FL-distal, 4 mice) and L5B PCs (n=8 neurons for sham and n=9 for FL, 4 mice). **(F)** Comparison of ERF and IRF between groups, obtained by counting total significant pixels in **A-D**. Note the L2/3 ERF in FL-distal was significantly increased compared to Sham and FL-proximal (unpaired t-test, P < 0.05); while in L5B, the IRF of both FL-proximal and FL-distal increased significantly (unpaired t-test, P < 0.05), instead of the ERF. **(G)** Laminal distribution of charge of EPSCs for L2/3 and L5B PCs in sham (black) vs. FL-proximal (blue) and FL-distal (red) groups. Left, L2/3 EPSCs strength in FL-distal group was significantly larger for infragranular layers compared to that in FL-proximal group (P<0.05). Right, no difference was found in EPSPs charge across layers between groups. **(H)** Same as **G** but for IPSCs. Left, L2/3 IPSCs charge was significantly lower in FL-distal group compared to that in FL-proximal group (P<0.05), mainly in supragranular layers. Right, L5 IPSCs strength was significantly lower in FL-distal group compared to that in FL-distal group (P=0.013), but mainly in infragranular layers.

In contrast, L5B PCs in proximal and distal groups both exhibited hypoexcitability (more obvious in proximal region), with the median E/I ratio values of 0.1 and 0.3, compared to the sham group (0.4, Figure S4D, *P*<0.01 and *P*<0.05, respectively). The IRF increased significantly in L5B PCs of both proximal and distal groups compared to the sham group (Figure 3F). As a result, the ERF size was larger than the IRF’s in the sham group (*P*<0.01), but in the FL group, the ERF size matched with IRF’s (Figure 3F, right panel). The combined results from both L2/3 and L5B data highlight the location-, layer- and synaptic origin-specific aberrant circuitry in FL cortices.

In all above experiments we did not separate the contribution from specific layers or input source to the E/I balance. To further examine this, we next used layer- and interneuron cell type-specific strategy to study the aberrant circuitry associated with FL-treatment.

### L4-derived synapses contribute to imbalanced E/I ratio in aberrant circuitry

L4 spiny neurons carry bottom-up information from the thalamus to cortical L2/3 neurons (Bureau et al., 2006; Lubke et al., 2003; Thomson and Bannister, 2003) and L5B PCs (Petreanu et al., 2009). Does maladaptive hyperexcitability (for L2/3 neurons) and hypoexcitability (for L5B PCs) result from differential re-organization of L4 synaptic input? We addressed this question by examining L4-specific inputs to L2/3 and L5B PCs in sham and FL mice. L4-specific expressions of ChR2 were achieved via injection of flexed ChR2 in Scnn1a-cre mice (see Methods, Figures 4A and 4D, (O’Connor et al., 2013)). Sham L2/3 PCs showed a centralized pattern in excitation maps and a bimodal distribution in inhibition maps (Figure 4B), presumably due to lateral inhibition from a neighboring barrel column. This lateral inhibition was also found in L5B PCs in the sham group (Figure 4C). L2/3 (Figure 4E), but not L5B PCs (Figure 4F), in FL groups displayed increased E/I ratios compared to their counterparts in the sham groups (Figure 4H, Figures S5A-5E, neuronal E/I ratio in L2/3:0.66 vs 0.37, P<0.05; L5B:0.22 vs 0.18, P>0.05). In addition, the lateral inhibition observed in sham PCs was interrupted in L2/3 (Figures 4E vs 4B) but not L5B PCs (Figures 4F vs 4C) in the FL group. The ERF size increased significantly in FL L2/3 PCs (Figure 4I, P<0.01), a change similar to the IRF size increase in FL L5B PCs (Figure 4I, P<0.001). These results suggest that the L4 → L2/3 synapse contributes to the reorganization of interlaminal and inter-columnal circuits.

**Figure 4.**
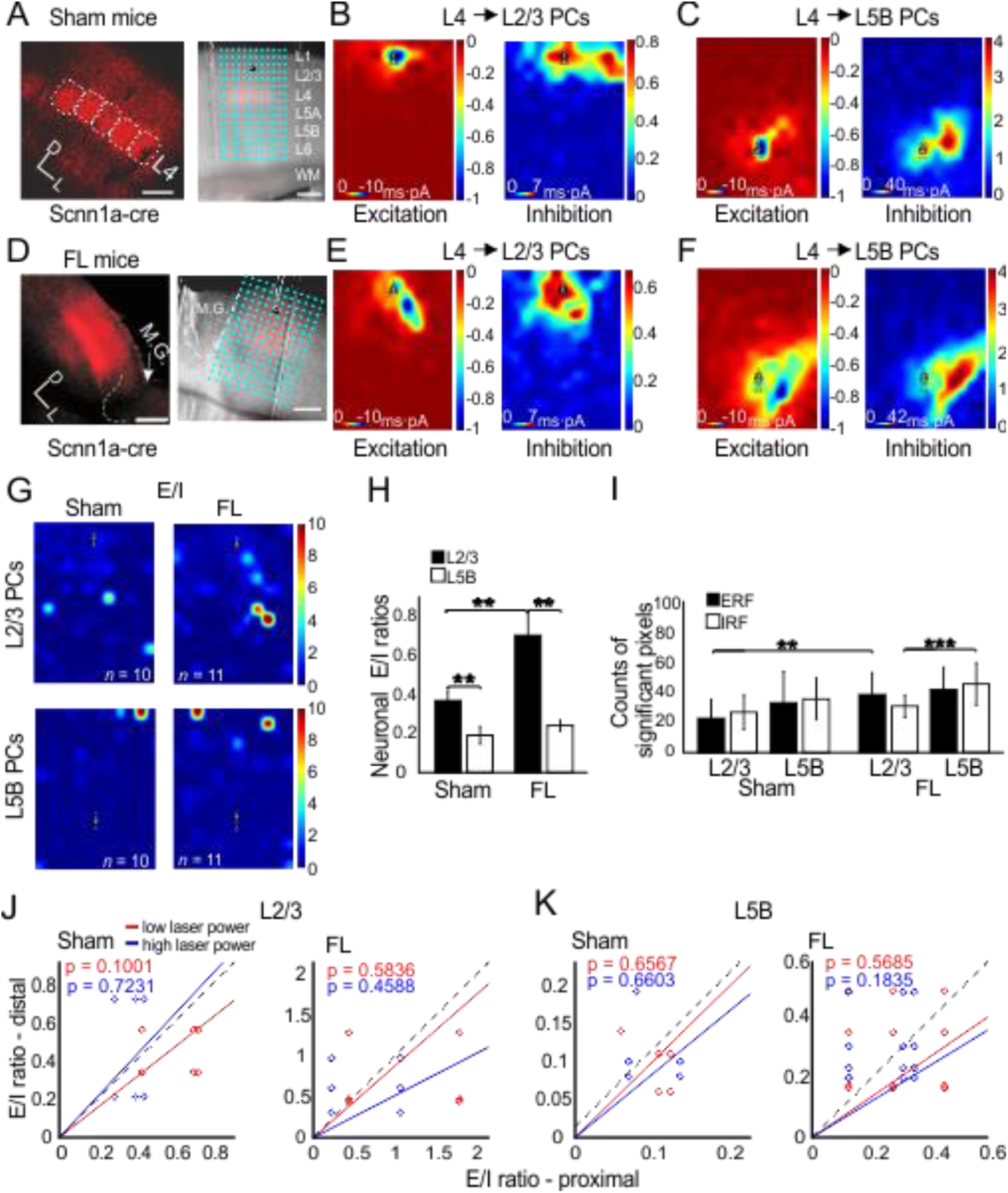
Input from granular layer contributes to the E/I imbalance in epileptic mice. **(A)** Left, confocal image shows the barrels in vS1 expressing ChR2 from a sham Scnn1a-cre mouse. Right, an exemplar recording from a L2/3 PC. Scale bar, 200 µm. (**B-C)** CRACM of synaptic input from L4 to L2/3 PCs (**B**) or to L5B PCs (**C**) in sham mice. Note the scale difference in Inhibition maps between two layers. **(D-F)** Same as **A-C** but for FL mice. The pixel number of inputs to L5B PCs for both Excitation and Inhibition increased (**F**). (**G**) E/I heat maps for L2/3 (Sham, n=10, FL, n=11) and L5 PCs (Sham, n=11, FL, n=11). **(H)** The averaged E/I ratios across neurons in FL were significantly higher in L2/3 PCs (P<0.01) but not in L5B PCs. **(I)** The ERF size in FL was also significantly higher in L2/3 (P<0.01); in contrast, the IRF, instead of ERF, increased significantly in L5B (P<0.001), but not L2/3. **(J)** The relationship between E/I ratios in Sham vs FL mice did not change under two different laser intensities.

To determine whether the circuit differences in proximal vs. distal regions were contributed by differential L4 inputs, we compared the proximal vs. distal neuronal E/I ratios in L4 → L2/3 and L4 → L5B PCs. The results showed the neuronal E/I ratios did not change significantly for either L4 → L2/3 or L4 → L5B PCs between proximal and distal groups (Figures 4J and 4K), suggesting the overabundant excitatory input to FL-distal L2/3 PCs observed previously came from sources other than local L4, probably from intra- and/or inter-columnar L5B (Figure 2C) and local L2/3 PCs. Together, these data suggest that the L4 synaptic inputs only partially contributed to the hyperexcitability in L2/3 PCs. However, the hypoexcitability in L5B PCs remains unclear. Is it due to altered inhibition from a specific subtype of interneuron? Is it a compensatory effect of cortical circuitry in response to epileptogenesis?

### Maladaptive organization of inhibitory circuits in FL cortex

We next examined if inhibitory synapses from distinct interneuron subtypes were maladaptively changed in L2/3 and L5B of the FL cortices. PV and SOM INs are the primary subtypes providing direct synaptic inhibition onto PCs (Rudy et al., 2011) and thus were the subjects of study here. PVs mainly target the perisomatic region of pyramidal neurons (Packer and Yuste, 2011), whereas SOMs innervate distal dendritic areas (Silberberg and Markram, 2007). We examined the direct monosynaptic input from PVs or SOMs onto L2/3 and L5B PCs (Figure 5, see Methods). As expected, CRACM revealed a perisomatic inhibition pattern in L2/3 and L5B PCs in PV-cre mice (Figure 5B and 5C). However, PV → L5B PCs in the FL group displayed a lower level of IPSC activities than the sham (Figure 5C, *P*<0.05), while PV → L2/3 PCs in the FL group did not. The IRF size of L5B PCs also dropped significantly (Figure 5E, P<0.05), presumably due to reduced synapses from PV INs. In SOM-cre mice, CRACM demonstrated an apparent dendritic targeting pattern of SOM input to PCs in both layers as expected (Figure 5G and 5H). While SOM → L2/3 PCs did not differ in their normalized pixel IPSC charge between the sham and FL (Figure 5I), SOM → L5B in the FL-treated mice showed much higher level of IPSC activities than in the sham-treated (Figure 5I, *P*<0.01). Meanwhile, the IRF size of L5B PCs in the FL group increased dramatically, suggesting more dendritic inhibitory synapses formed by SOM in FL-treated groups (Figure 5J). This demonstrates that SOM, but not PV, contribute to the hypoexcitability in L5B PCs.

**Figure 5.**
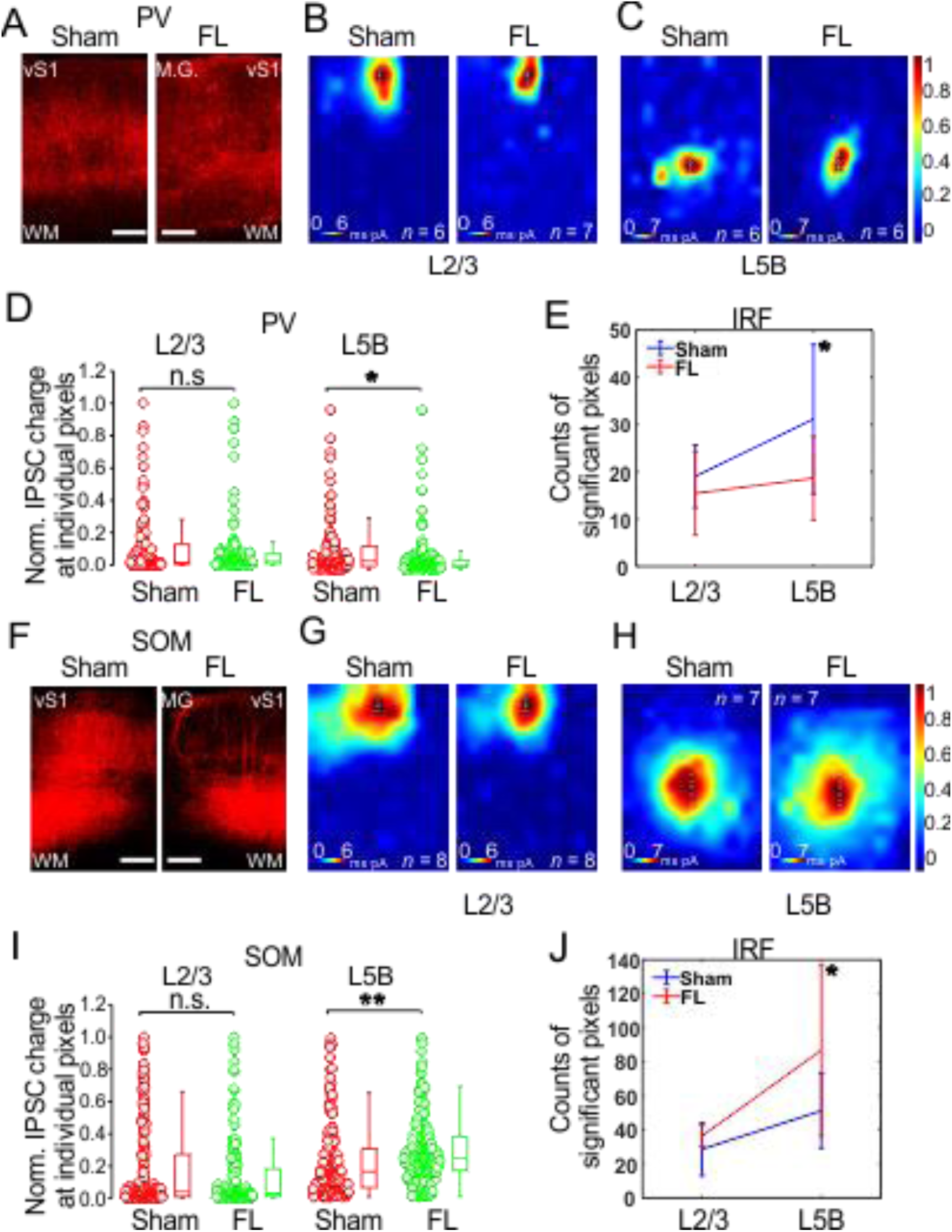
Inhibitory inputs from PVs and SOMs to PCs of vS1 in epileptic mice. **(A)** Left, confocal images show ChR2 expression in PVs for both sham and FL slices. **(B)** Heat maps show the inhibitory input from local PVs to L2/3 PCs. **(C)** Heat maps show the inhibitory input from local PVs to L5B PCs. **(D)** The normalized IPSC charge at individual pixels across neurons did not differ in L2/3 PCs between sham and FL. However, it was significantly reduced in L5B for FL group (P<0.05), suggesting PV functional output was impaired in infragranular layer. **(E)** The IRF in L5B was also significantly reduced in FL group (sham, n=6, 3 mice; FL, n=6, 3 mice P<0.05). **(F)** Left, confocal images show ChR2 expression in SOMs for both sham and FL slices. **(G)** Heat maps show the inhibitory input from local SOMs to L2/3 PCs. **(H)** Heat maps show the inhibitory input from local SOMs to L5B PCs. **(I)** The normalized IPSC charge at individual pixels across neurons did not differ in L2/3 PCs between sham and FL. However, it was significantly increased in L5B for FL group (P<0.01), opposite to the effect of FL on PV INs. **(J)** IRF in L5B was also significantly increased in FL group (sham, n=7; FL, n=7, P<0.05).

We also compared the absolute IPSC charge between proximal and distal regions for both sham and FL groups (Figure S6). As expected, the IPSC charge of the sham PV → L2/3 PCs did not differ between the two regions (Figure S6A, *n* = 12 pairs). However, FL PV-L2/3 PCs showed significantly lower IPSC charge in FL-distal compared to FL-proximal region (Figure S6A, *P*<0.05, *n* = 12 pairs). Surprisingly, SOM → L2/3 PCs in FL-distal regions showed significantly lower IPSC charge than in FL-proximal regions (Figure S6C, *P*<0.001, *n* = 16 pairs). The L5B IPSC charge of FL-proximal vs FL-distal regions did not show a significant difference for both PV and SOM INs (Figures S6B and 6D). These results suggest the functions of both PV and SOM INs in L2/3 were compromised in distal FL region, leading to the hyperexcitable states in L2/3 of this region. The hypoexcitable state in L5B was primarily caused by enhanced SOM inhibitory innervation of the distal dendritic sites of L5B PCs, which is presumably caused by axonal sprouting of SOM INs (Figure 5H). If this is the case, we predict that optogenetic manipulation of these SOM inputs will have larger influence over the chronic spike-wave seizures.

### Closed-loop optogenetic activation of remaining SOMs, but not PVs, distal to microgyrus irreversibly abolished spontaneous SWDs *in vivo*

To further examine if SOM or PV INs responded to epileptic states as a compensatory mechanism, we applied closed-loop optogenetic stimulation of both types to test the effect *in vivo*. Flexed ChR2 was injected in FL-distal regions in PV- and SOM-cre mice, respectively, and a fiber-optic probe was implanted in same regions (Figure 6A and 6D). After waiting for at least two weeks upon viral expressions, we performed EEG recordings and closed-loop optogenetic stimulations. Once SWD activities were detected (Figure 6B and 6E, based on the characteristic α (8-12 Hz) and θ (4-8 Hz) activity of the SWD (Williams et al., 2016)), we applied closed-loop optogenetic activation (5Hz) of PVs or SOMs in distal regions of microgyrus. Our results showed that optogenetic activation of SOM, but not PV INs, irreversibly abolished the spontaneous SWD events (Figure 6C and z6F), suggesting that enhancing remaining dendritic inputs from SOM INs, but not the perisomatic inhibition from PV INs, eliminated the characteristic α (8-12 Hz) and θ (4-8 Hz) activities of the SWD in FL-treated mice..

**Figure 6.**
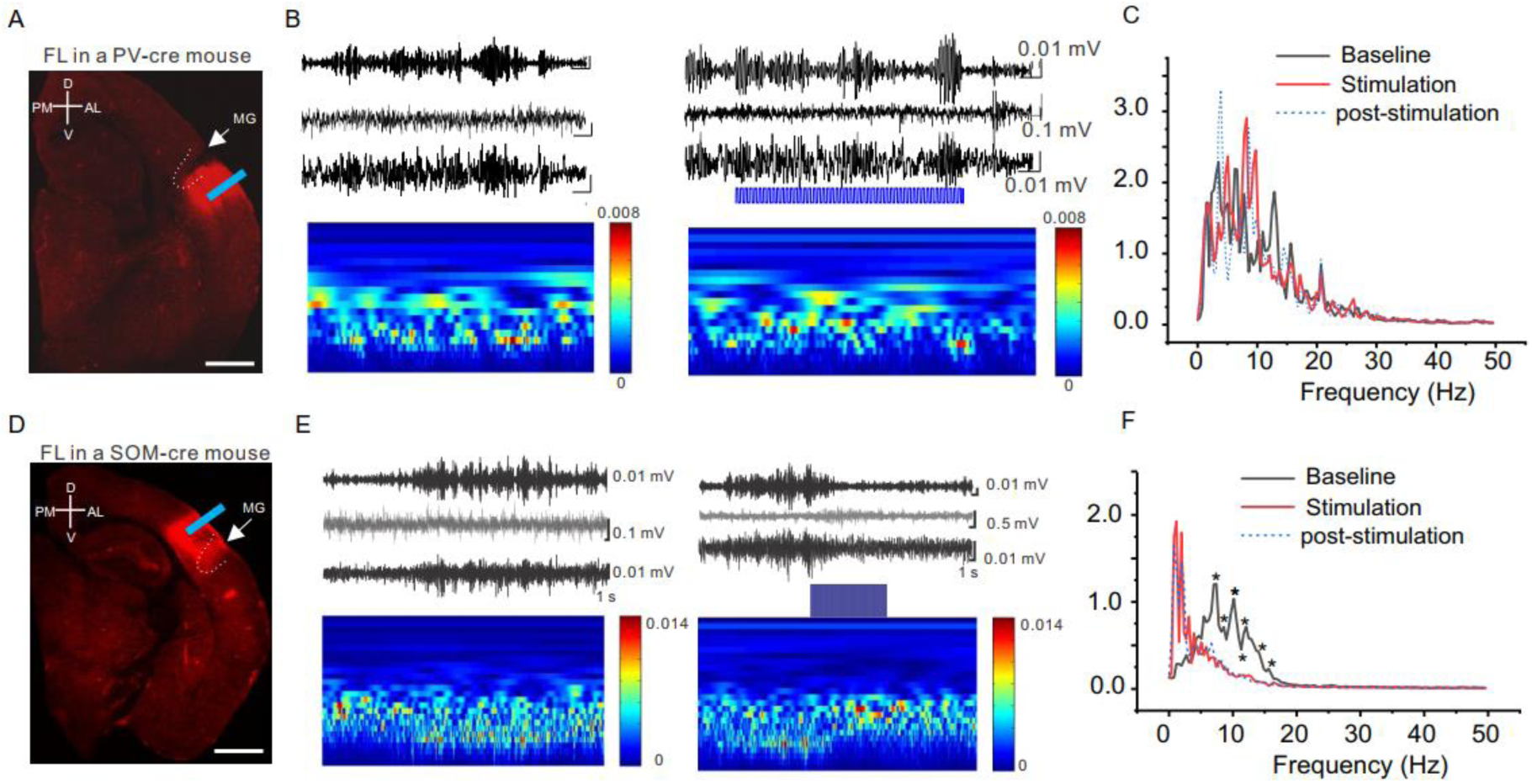
Close-loop optogenetic activation of SOM but not PVs terminated the spontaneous SWD *in vivo*. **(A)** Confocal image show ChR2 expression in a FL PV-cre mouse. **(B)** EEG traces (top) and time frequency plot (continuous wavelet transformation of the same EEG traces) show SWD before, during and after optogenetic stimulation (5Hz) in the peri-microgyri area. **(C)** Power spectrum of the baseline, optogenetic stimulation, and post-stimulation period (2 seconds in each period, n=8 mice). **(D-F)** Same as **A-C** but for SOM-cre mice. Note the power was significantly reduced at the frequency ranging from 7-15 Hz upon optical stimulation compared to baseline (n=5 mice, P<0.05).

## Discussion

Developmental disruption of neuronal migration underlies the formation of cortical microgyrus. Chronic spontaneous electrographic seizures are a core feature of our focal MCD model. We used *in vivo* and *in vitro* electrophysiology together with laser scanning photostimulation mapping, to characterize how aberrant synapses lead to epileptic circuitry. Our data suggests that excessive synaptic input from granular and infragranular layers are major driving forces to the hyperexcitability of supragranular PCs. In sharp contrast, the otherwise active L5 PCs experienced hypoexcitability in a FL cortex. Furthermore, we found that the chronic hyperexcitability foci is located at regions distal to the microgyrus, both *in vivo* and in *vitro*. Strikingly, despite of functional deficits of both PVs and SOMs in the infragranular layer, their postsynaptic innervation sites differed significantly, as IRF size dropped for PV but increased for SOM. Neither functional deficit nor RF change has been found for either subtype of interneurons in L2/3. Importantly, activation of SOMs at FL-distal regions abolished epileptic activities irreversibly, at least during the time period we tested.

It has been well documented that synaptic excitation and inhibition in cortical neurons shows a proportionality, or balance, through the use of recurrent local synaptic connections to maintain normal brain functions (Bridi et al., 2019; Isaacson and Scanziani, 2011; Okun et al., 2015; Shu et al., 2003; Xue et al., 2014; Yang and Sun, 2018; Zhang et al., 2011; Zhou et al., 2014). L2/3 and L5B PCs play different roles in the horizontal propagation of recurrent-network activities, including epileptiform activities (Telfeian and Connors, 1998; Wester and Contreras, 2013). In canonical cortical circuitry, L2/3 receives its prominent excitatory drive from L4 and subsequently sends major output to L5B (Thomson and Bannister, 2003). In a healthy barrel cortex, L2/3 PCs display a lower synaptic E/I ratio compared to L5B PCs within same columns (Adesnik and Scanziani, 2010; Yang and Sun, 2018; Zhang et al., 2011), consistent with their respective functions, e.g. the sparse coding strategy used by supragranular neurons for feature integration (Crochet et al., 2011; Harris and Mrsic-Flogel, 2013) and active feedback to subcortical regions (Groh et al., 2010; Li et al., 2015). L2/3 PCs also send their output to other ipsi- and contra-lateral cortical regions. Their hyperexcitability in the malformed cortex might facilitate abnormal intracortical propagation and the generalization of epileptiform activities. Intrinsically bursting L5B PCs have also been implicated in the generation of epileptiform activities due to their intrinsic properties, extensive horizontal intracortical axons, and the feedback modulation of ascending sensory information (Vogels et al., 2011; Xue et al., 2014). However, our results indicate that circuits formed by L5B PC neurons are hypoexcitable in FL vS1. This result is consistent with our *in vivo* extracellular data, which shows an absence of optogenetically induced epileptiform activities. However, the hypoexcitability in L5B PCs does not mean that they were not involved in the induction of hyperexcitability in L2/3 PCs. On the contrary, our *in vivo* data regarding latency of epileptiform bursts (Figure 2C) indicates that spontaneously occurring hyperexcitable activities in L2/3 at least partially originated from deeper layers. Thus, our results lead to a new hypothesis that the interplay between infra- and supra-granular layers is necessary for the chronic epileptogenesis, consistent with an idea put forward by previous studies (Telfeian and Connors, 1998; Wester and Contreras, 2012), regarding the propagation of self-sustained activities in a normal neocortex. This hypothesis also suggests that the normal long-range neocortical network has been hijacked to give rise to epileptiform activities.

In an epileptic brain, the E/I imbalance directly leads to epileptic discharges of the “epileptic neuron” (Prince and Wilder, 1967). However, the mechanisms contributing to epileptogenesis in malformed cortices are largely unknown. It is also unknown why seizures associated with cortical malformations are medically intractable (Schwartzkroin and Walsh, 2000; Wong, 2008). A focal epileptogenesis mechanism has been proposed in previous studies, characterized by an initiation of epileptiform activities in an adjacent 6-layered cortical area distal to the microgyrus (Jacobs et al., 1996; Luhmann, 2016). This hypothesis has yet to be tested in a mouse model exhibiting spontaneous seizures. Importantly, severing a microgyrus from an adjacent cortical region does not prevent the initiation of epileptiform activities (Jacobs et al., 1996; Luhmann, 2016). Human MCD is rarely a localized, or even regional process, but is instead an electrophysiologically, functionally, and ultimately clinically integrated neural network disorder. Our results demonstrate the aberrant ectopic inter-laminar synaptic inputs (primarily from L5B) to L2/3 neurons in distal paramicrogyral areas may be a critical constituent of epileptogenesis. This epileptic tendency could be further shaped by differential compensatory effects in subtypes of INs. This difference was not due to IN density change in the FL model (Hablitz and DeFazio, 1998; Schwarz et al., 2000). The hypoexcitability of L5B PCs could be partially explained by axonal sprouting of SOM INs within the same layer(Jiang et al., 2018), indicating a selected disturbance of IN subtypes in this model. This aberrant elevated activity of SOM INs might counteract the excitatory drive from hyperexcitable L2/3 PCs, which might explain how closed-loop activation of SOMs alleviates epileptogenetic activities. Other untested possibilities include excessive excitatory drive in INs as well as drive from intralaminar input or thalamocortical input (Meeren et al., 2009). GABA_A_ receptor reduction and increased AMPA receptor density in the paramicrogyral area, as reported earlier (Zilles et al., 1998), might also contribute to imbalanced E/I ratio. Understanding the fine layer- and cell type-specific circuitry that gives rise to aberrant activities in the epileptic brain is an important step towards a revolutionized novel treatment.

## Acknowledgements

Funding for this research provided from NINDS/NIH grant 5R01NS094550 and P20GM121310. We thank Sun lab members for assistance with animal care and breeding, proof reading of the manuscript and Chen Zhou for performing the EEG electrode implants and recordings and closed-loop optogenetic stimulations.

## Author Contributions

W.Y. and Q.Q.S. conceived the project. W.Y. performed the CRACM experiments. A.W. performed the in vivo electrophysiology experiments. W.Y. and A.W. analyzed the data. W.Y. and Q.Q.S. wrote the paper with help from A.W.

## Declaration of Interests

None of the authors has any conflict of interest to disclose.

## Star*Methods

Detailed methods are provided in the online version of this paper and include the following:

- KEY RESOURCES TABLE
- LEAD CONTACT AND MATERIALS AVAILABILITY
- EXPERIMENTAL MODEL AND SUBJECT DETAILS
- METHOD DETAILS
  Experimental microgyrus in mice
  Stereotaxic viral injection
  Microelectrode arrays
  In vivo Extracellular recordings
  Intracranial EEG (iEEG) recordings
  Closed-loop optogenetic stimulation
  Histology after in vivo recording
  Data and statistical analysis for in vivo extracellular recordings
  Slice preparation and electrophysiology
  Photostimulation and CRACM
  LED field stimulation
  Data analysis for slice mapping experiments
- QUANTIFICATION AND STATISTICAL ANALYSIS
- DATA AND CODE AVAILABILITY
- KEY RESOURCES TABLE

Unique link: https://star-methods.com/?rid=KRT5e9decf29da8b

- LEAD CONTACT AND MATERIALS AVAILABILITY

Further information and requests for resources and reagents may be directed to and will be fulfilled by Lead Contact, Dr. Qian-Quan Sun. This study did not generate new unique reagents.

- EXPERIMENTAL MODEL AND SUBJECT DETAILS

All experiments were performed under protocols approved by the Institutional Animal Care and Use Committee of the University of Wyoming. Mice were housed in a vivarium maintained at 22-23 ⁰C on a 12:12 h light-dark cycle. Food and water were available *ad libitum*. Mice of both sexes were used in this study. Unless otherwise specified, mice of a CD-1 background were used. Scnn1a-cre mice (Jackson lab) were used for L4-specific experiments; PV-Cre (Jackson lab) and Som-Cre (Jackson lab) mice were used for inhibitory CRACM experiments. All cre lines were of a C57BL/6J background. Both C57BL/6J and CD-1 mouse strains had been used for an earlier study of FL induced spontaneous spike-wave seizures and no significant differences were found between the two strains (Williams et al., 2016).

## Methods Details

### Experimental microgyrus in mice

An experimental microgyrus was induced in postnatal day 0 (P0) mouse pups via the use of the transcranial freeze-lesion (FL) as previously described (Dvorak and Feit, 1977, 1978; Dvorak et al., 1978; Jacobs et al., 1999; Sun et al., 2016; Williams et al., 2016). The pups were first briefly anesthetized by surrounding them with ice for ~2 minutes, the skull was exposed, and a copper bar (1 mm tip diameter and pre-submerged in liquid nitrogen for ~10-min) was gently placed on the surface of right barrel cortex region for ~2 s (from bregma: 2.2±0.2 mm lateral and 0±0.1 mm AP). Controls were sham-operated littermates treated the same as FL pups except for the use of another copper bar kept at room temperature (23 °C). The scalp was sutured with super glue. Treated mice were allowed to recover for ~30-min in a heated cage before being returning to the dam. Examination of slices for electrophysiological experiments at age of P46-180 or post hoc histological observation indicated 4-layered abnormal cortical structures were reliably induced as previously reported (Dvorak and Feit, 1977, 1978; Dvorak et al., 1978; Jacobs et al., 1999; Williams et al., 2016) (Figure S2C). The microgyrus was mainly located around central or lateral side of the barrel field. The recorded neurons were most located medial to the FL-induced microsulcus. Some neurons were located lateral to the microsulcus, which showed similar results to the medial neurons, and thus were included for analysis. The microgyrus with an abnormal 4-layered structure typically spanned a 400 ± 80 µm (tangential) region (Figure S2C). Cortical regions surrounding the microgyrus displayed normal 6-layered structure as previously reported (Dvorak and Feit, 1977, 1978; Dvorak et al., 1978; Jacobs et al., 1999; Williams et al., 2016). We defined a region 400-1000 µm away from center of FL-induced microsulcus as “proximal” and that >1000 µm as “distal” (<2000 µm). While most recorded neurons were located in S1 barrel cortex, some neurons in distal regions fell into the S1 trunk region.

### Stereotaxic viral injection

Virus injection was performed in sham or FL mice aged P14-P16. Mice were anesthetized with 3% isoflurane (vol/vol) and maintained with oxygenated 2% isoflurane mixture throughout the surgery procedure. AAV2.1.CAG.hChR2(H134R)-mCherry (University of Pennsylvania Vector Core) was injected in vS1 area of CD1 mice. AAV2/1.CAGGS.flex.ChR2.tdTomato.WPRE.SV40 (University of Pennsylvania Vector Core) was injected in vS1 of Scnn1a-cre mice, PV-cre mice, or SOM-cre mice. The viral vector (2 µl) was loaded into the tip (~10 µm in diameter) of a beveled glass micropipette (Drummond Scientific Co.). A custom stereotactic apparatus was used to deliver the viral vector to the cortex through a small hole drilled in the skull. For vS1 injection, the virus was injected at two depths: 400 µm and 800 µm; for L4 injection, 400 µm and 600 µm. For each depth, a volume of ~100 nl was injected within 1-2 minutes using a micromanipulator (MP-285-system, Sutter Instrument). Injection pipette was kept in place for 5 minutes for each depth after injection. Injected mice were warmed in a heated cage until full recovery from surgery before being returned to the dams. After weaning day, pups were separated by gender until experiments. Electrophysiological experiments would not start until at least two weeks after viral injection.

### Microelectrode arrays

Extracellular signals were recorded using linear microelectrode SmartProbe^TM^ arrays (NeuroNexus, Ann Arbor, MI) from the vS1 of control and FL mice (Williams and Sun, 2019; Williams et al., 2016). The electrode array consisted of a single shank with 16 individual electrodes separated by 100 μm, with on-board electronics for digital conversion of the signals, and linked to a SmartBox^TM^ control and data streaming system through SmartLink headstage (NeuroNexus). The array was positioned perpendicular to the pia. Each signal was filtered digitally (1-10000 Hz band-pass and 60 Hz notch filters) and recorded with a sample rate of 20 kHz using Smartbox 2.01 (NeuroNexus). Additional off-line digital filtering was used to define local field potentials (LFPs, 1-100 Hz) and multi-unit (MU) activity (800-5000 Hz).

### In vivo Extracellular recordings

All extracellular recordings were obtained from anesthetized (0.5-1.0% isoflurane delivered in oxygen) and head-restrained mice (stereotaxic frame supplied with mouse and gas anesthesia adaptors, Stoelting, Wood Dale, IL) as previously described (Williams et al., 2016). Normal animal body temperature was maintained during recordings with a circulating water bath (Haake, Thermo Electron, Newington, NH) infused custom heating pad and continuous rectal temperature probe (BK precision, Yorba Linda, CA). A scalp incision was made to expose the skull over the right vS1. The recording stage was performed in a custom-built Faraday cage on vibration isolation table (Vibraplane, Kinetic Systems, Boston, MA). Single or multiple burr holes (1 mm diameter) were made through the skull with a dental drill for stereotaxic insertion of linear microelectrode arrays and fiber optic probes (laser stimulation). Fiber optic probes were placed directly over the virus injection ports. The microelectrode arrays were positioned laterally within 1 mm of the fiber optic probe at an angle of 40 degrees from vertical for perpendicular insertion into the S1BF. A 3-axis micromanipulator (hydraulic, Narishige, Amityville, NY) was used to advance the array slowly into the brain and was then waited for 20 minutes once in place (~1650 μm depth). Electrode placement was visually guided under a dissecting microscope to verify that all electrodes had penetrated the surface of the brain. The abrupt change in the local field potential (LFP) between cortical and subcortical layers was used as a visual marker to verify electrode depth (Williams and Sun, 2019; Williams et al., 2016). The first 12 electrodes were subsequently analyzed for laser-evoked multi-unit (MU) activity. The dura was found to offer only minimal resistance to electrode insertion and was therefore left intact. Warm physiologically isotonic saline was administered to the craniotomy site during recording period. The silver wire reference electrode was placed under the skin flap of the scalp incision.

### Intracranial EEG (iEEG) recordings

Methods of iEEG surgery, recording and analysis were similar to the methods described in our previous publication (Sun et al., 2016; Williams and Sun, 2019; Williams et al., 2016). Briefly, polyimide-insulated stainless steel wires (125 µm, California Fine Wire) and connecting pins will be implanted into S1 (right side S1R), VPM and contralateral hippocampus. Upon animals fully recovering from the surgery, EEG recordings will be performed in a 24-hour cycle with simultaneous video behavior monitoring and automated infrared (IR)-activity tracking at a frequency of once per week. EEG signals will be amplified via a differential AC amplifier (Model 1700, A-M system), digitized at 1000Hz using Power 1401, and analyzed using the Spike-2® program.

### Closed-loop optogenetic stimulation

We implanted a multi-mode fiber optic patch cable (Ø150µm, Thorlabs) via a 1.25mm OD multimode ceramic zirconia ferrule (Precision Fiber Products, Inc, Milpitas, CA 95035), which was glued together with the S1R electrode to form an optrode configuration and implanted in the distal site (~1-1.5mm) of the microgyrus location (Fig 6a, d). The multi-mode fiber optic patch cable was coupled to a blue laser (473nm, 100 mW, DPSS blue laser, www.lasercentury.com) via a SMA mating end. Laser pulses were delivered via a custom made pulse generator at three frequencies (1, 5 and 10Hz). It was estimated that an area of approximately 0.3mm depth and 0.3mm width was illuminated upon activation of the fiber based on tests using similar fiber in brain slice *in vitro* (Wang and Sun, 2012). During the online recording period, bandwidth filtered (8-15Hz) of the EEG trace is displayed for the detection of spike-wave seizures, which is defined based on our initial characterization(Sun et al., 2016). Upon detection of spike-wave seizures, laser stimulation is delivered for a period of 5-10 seconds.

### Histology after *in vivo* recording

Following all in vivo electrophysiology recordings, animals were anesthetized and transcardially perfused with 0.9% saline followed by 4% paraformaldehyde. Brains were extracted, immersed in 4% paraformaldehyde for 24 h, and then transferred to 0.1 M phosphate buffer containing sequentially increasing levels of sucrose (10%, 20%, 30%, pH 7.4, 4°C) across 3 consecutive days. Brain tissue was then removed from the sucrose solution and cut into serial sections through the area of interest. The microelectrode was immersed with DiI (2 mg/ml in ethanol,). Several techniques were used to visualize the probe track in relation to the freeze lesion and S1 barrel field. Thick coronal sections (300 μm) were cut on a vibratome for direct visualization of the probe track (DiI) in relation to the freeze lesion. A cryostat was used to cut thinner coronal sections (50-80 μm) for staining with cytochrome oxidase (CO) to visualize the barrel cortex in relation to the freeze lesion. Coronal sections were incubated at room temperature in a solution of CO (0.5 mg/ml, Sigma), sucrose (0.4 mg/ml, Sigma), and DAB (0.625 mg/ml, Sigma) in 0.1 M phosphate buffer (pH 7.4) for 60-90 min until cortical barrels were visible under a dissecting microscope. All sections were then mounted on glass slides with a DAPI mounting medium (Vectashield, Vector Laboratories, Burlingame, CA) and coverslipped. Brain images were evaluated under a light/fluorescent microscope (Zeiss Axioskop 2, Ontario, CA) and digitally imaged using Axiovision software (ver. 4.6, Zeiss). Histological analysis of serial brain sections was used to construct cortical maps to mark the location of the recording sites in relation to the induced microgyric lesions.

### Data and statistical analysis for *in vivo* extracellular recordings

NeuroExplorer (Nex Technologies, Madison, AL) was used for data analysis, visual inspection, and digital filtering of MU activity. Power spectral density (PSD) values were computed across a frequency range of 1-20 Hz with a single taper Hann FFT using a bin size of. 0012 Hz and 50% overlap between bins to mitigate data loss at the spectral edge of each bin. Correlation analysis was used to compute the cross-correlation and latency shift between simultaneously recorded signals from a single linear microarray recording using standard correlograms across a −0.3 to +0.3 sec offset (bin size of 0.05 ms based on the A/D sampling rate)(Williams and Sun, 2019; Williams et al., 2016). Data files were exported to Spike2 (Cambridge Electronic Design Limited, Cambridge, England) for spike sorting. MU spikes were detected using template matching software triggered by negative threshold deflections greater than −0.02 mV (Spike2 software, see Figures 2D-2F) within a 50 ms window surrounding each laser pulse (10 ms pre-stimulus recording). Post-stimulus time histograms and raster plots of MU activity were computed from 32 laser-evoked responses at each recording site (NeuroExplorer) and spike count averaged across each group (Sigmaplot). Following analysis, raw data was exported to Microsoft Excel for tabulation of statistical averages and standard error values or Sigmaplot (ver. 11.0, SYSTAT software Inc., San Jose, CA) for graphical display and statistical analysis between groups. Group averages are presented mean ± S.E.M. A multifactorial analysis of variance (ANOVA) was used to evaluate main effects and interactions when multiple independent variables were present including spike count, electrode, laser intensity, time, experimental group (e.g. control vs. FL), followed by a Holm-Sidak post-hoc analysis to evaluate significant differences between individual groups. A P value of < 0.05 was considered significant.

### Slice preparation and electrophysiology

Mice aged P46-180 were decapitated under isoflurane anesthesia. The brain was quickly removed and placed into ice cold cutting solution (in mM: 2.5 KCl, 1.25 NaH_2_PO_4_, 10.0 MgCl_2_, 0.5 CaCl_2_, 26.0 NaHCO_3_, 11.0 glucose and 234.0 sucrose). Coronal brain slices were cut (TPI, St. Louis, MO) to a 300-µm thickness. Slices were incubated in 35°C oxygenated (95% O_2_ and 5% CO_2_) aCSF (in mM: 126.0 NaCl, 2.5 KCl, 1.25 NaH_2_PO_4_, 1.0 MgCl_2_, 2.0 CaCl_2_, 26.0 NaHCO_3_ and 10.0 glucose, 295 mOsm) for 1 h. The incubation chamber was then kept at room temperature for up to 5 h throughout experiments. All recordings were performed at room temperature in circulating aCSF (1ml/min). Only slices with prominent barrels were used. Whole-cell recordings were obtained using borosilicate glass pipettes with a filament (resistance 4-5 MΩ). Recording pipettes were filled with a Cs-based intracellular solution (in mM: 120 cesium gluconate, 10 phosphocreatine-Tris, 3 MgCl_2_, 0.07 CaCl_2_, 4 EGTA, 10 HEPES, 4 Na_2_-ATP, and 1 Na-GTP) with the pH adjusted to 7.35 with CsOH (289 mOsm). Neurobiotin was always included in the intracellular solution (0.5%, wt/vol, Vector Labs). Whole-cell recordings were obtained with an Axopatch 700B amplifier (Axon Instruments) and a 1322A. Data from CRACM experiments were collected using the Matlab-based software *Ephus (Suter et al., 2010)*. Cells were recorded at depths from 50 to 110 µm within the brain slice. For CRACM experiments, EPSCs were recorded via a voltage-clamp with holding potential at −45 to −48 mV (L2/3 PCs) or −45 to −53 mV (L5B PCs) such that a fast (several ms) downward current was evoked when a laser beam was directed to recorded neuronal soma position. IPSCs were recorded in the same mode with holding potential at 0 mV. L2/3 and L5B PCs were verified by post hoc morphological confirmation of the existence of thick proximal dendrites.

### Photostimulation and CRACM

The blue laser beam (Shanghai Laser and Optics Century Co., 473 nm) was carefully aligned with routing mirrors, Pockels cell (ConOptics) and a pair of galvanometer scanners (Cambridge Scanning, Inc.) to generate a relatively cylindrical beam through a 4x objective (0.10 NA, Olympus) at the specimen plane (~15 µm in diameter, full-width at half max) as reported(Cruikshank et al., 2007; Wehr and Zador, 2003). A portion of laser beam was redirected to a photodiode (silicon detector, Edmund Optics) through a beam splitter (30:70 split, 450-650 nm range, Thorlabs). The detected voltage was coupled to laser intensity measurements using a handheld laser power meter (Edmund Optics) placed at the immediate back of the 4x objective. The majority of recorded neurons was measured under two different intensities. The laser power (0.1-1.6 mW, 1 ms duration) was adjusted such that the largest IPSC_CRACM_ had peak values ranging between 80-150 pA at lower intensity, while at stronger intensity the values ranged between 200-400 pA (Mao et al., 2011; Petreanu et al., 2007; Petreanu et al., 2009). Slices with insufficient virus expression (weak fluorescence) were excluded from subsequent analysis. Each cell was mapped with a 16 x 12 photostimulation grid (distance between adjacent points, 75 µm) covering the majority, if not all, of the polysynaptic inputs to dendritic arbor of recorded neurons. Each map was averaged from simulations repeated 3 times. The photostimulation sequence was given in a pseudo-random manner to maximize the intervals between adjacent locations receiving the photostimulation (Mao et al., 2011; Petreanu et al., 2007; Petreanu et al., 2009).

### LED field stimulation

In a subset of experiments, blue LED (LEDD1B, 488 nm, Thorlabs) field stimulation was used via a 4x objective to examine E/I ratio differences compared to CRACM (Figure S3). The light was pulsed for a duration of 2 ms. Light intensity (0.1-0.8 mW) was adjusted accordingly as mentioned above for a two-intensity stimulation. The interval between sweeps (20 repetitions) were 10 s. Holding potentials for the voltage clamp were identical to those used for CRACM experiments for same recorded neurons.

### Data analysis for slice mapping experiments

We performed data analysis using custom-developed software (MATLAB, MathWorks). Each trace of raw data in *Ephus* represented a 400-ms time window of averaged recorded current under the voltage clamp: the first 100 ms was the baseline before photostimulation, laser delivery at 100 ms time point and the following 300 ms showed the evoked EPSC/IPSC (Figure S1). The area of the postsynaptic potential defined as the integral of the recorded potential above baseline between 0 and 200 ms after laser onset was calculated. The pixel values of CRACM map corresponded to the average EPSC or IPSC charge calculated as above. This time window essentially captured the monosynaptic and major components of polysynaptic activities of evoked EPSC/IPSC. EPSC/IPSC charge within this time window was used to calculate their corresponding strength and ratios. For each pixel of a CRACM map (16×12=192 pixels in total), we calculated the charges for both EPSCs and IPSCs and converted them to heat maps (Figure S1). Within each averaged EPSC map and its corresponding IPSC map of each cell, the values of individual pixels were normalized to the pixel value with maximal charge within each EPSC map (e.g. Figure1A, two outside scale bars). Considering the variability of ChR2 expression across different animals, we further unified all the EPSC map scales of each class to the scale of sham-L2/3 experiments (e.g. Figure 1, two inside scale bars) by dividing or multiplying by a factor such that the maximum charge of each class of experiment was same. The scales of corresponding IPSC maps were also changed accordingly.

E/I ratio maps were derived by dividing the EPSC charge by IPSC charge (pixel E/I ratio, e.g. Figure 1E, pixels with only either EPSC or IPSC or neither were excluded). This analysis gave rise to a spatial distribution pattern of E/I ratios of one recorded neuron across its dendritic arbor or soma with polysynaptic input. The ratio limit was arbitrarily set as 10 (E/I ratio>10 was forced to 10). The E/I ratio is calculated from dividing the sum of individual pixel EPSC charge by the sum of individual pixel IPSC charge. This calculation generated one single value reflecting overall excitability profile of recorded neurons. Besides E/I ratio, response size for each synaptic class was also calculated to determine the polysynaptic or monosynaptic (for PV or SOM INs only) “receptive field” (RF) of each recorded neuron. Only pixels with significant responses (>3x standard deviation of the baseline) were included in analysis.

A paired *t* test was used for neuronal E/I ratio comparisons at the two intensities. Nonparametric Wilcoxon signed-rank tests were used for pixel size comparisons between paired groups as well as E/I ratio comparisons. Multivariate ANOVA was used for spectrum power analysis. One-way ANOVA was used for strength comparison between sham, FL-proximal and FL-distal groups (Figure 3G and 3H).

## QUANTIFICATION AND STATISTICAL ANALYSIS

Group averages are presented mean ± S.E.M. and a P value of < 0.05 was considered significant. Following analysis, raw data was exported to Microsoft Excel for tabulation of statistical averages and standard error values or Sigmaplot (ver. 11.0, SYSTAT software Inc., San Jose, CA) for graphical display and statistical analysis between groups. A multifactorial analysis of variance (ANOVA) was used to evaluate main effects and interactions when multiple independent variables were present including spike count, electrode, laser intensity, time, experimental group (e.g. control vs. FL), followed by a Holm-Sidak post-hoc analysis to evaluate significant differences between individual groups. A paired *t* test was used for neuronal E/I ratio comparisons at the two intensities. Nonparametric Wilcoxon signed-rank tests were used for pixel size comparisons between paired groups as well as E/I ratio comparisons. Multivariate ANOVA was used for spectrum power analysis. One-way ANOVA was used for strength comparison between sham, FL-proximal and FL-distal groups. Statistical methods were not used to pre-determine sample size.

## DATA AND CODE AVAILABILITY

The published article includes all datasets generated or analyzed during this study. Detailed datasets and codes supporting the current study are available from the corresponding authors on request.

## Supplemental Figures and Legends

**Figure S1.**
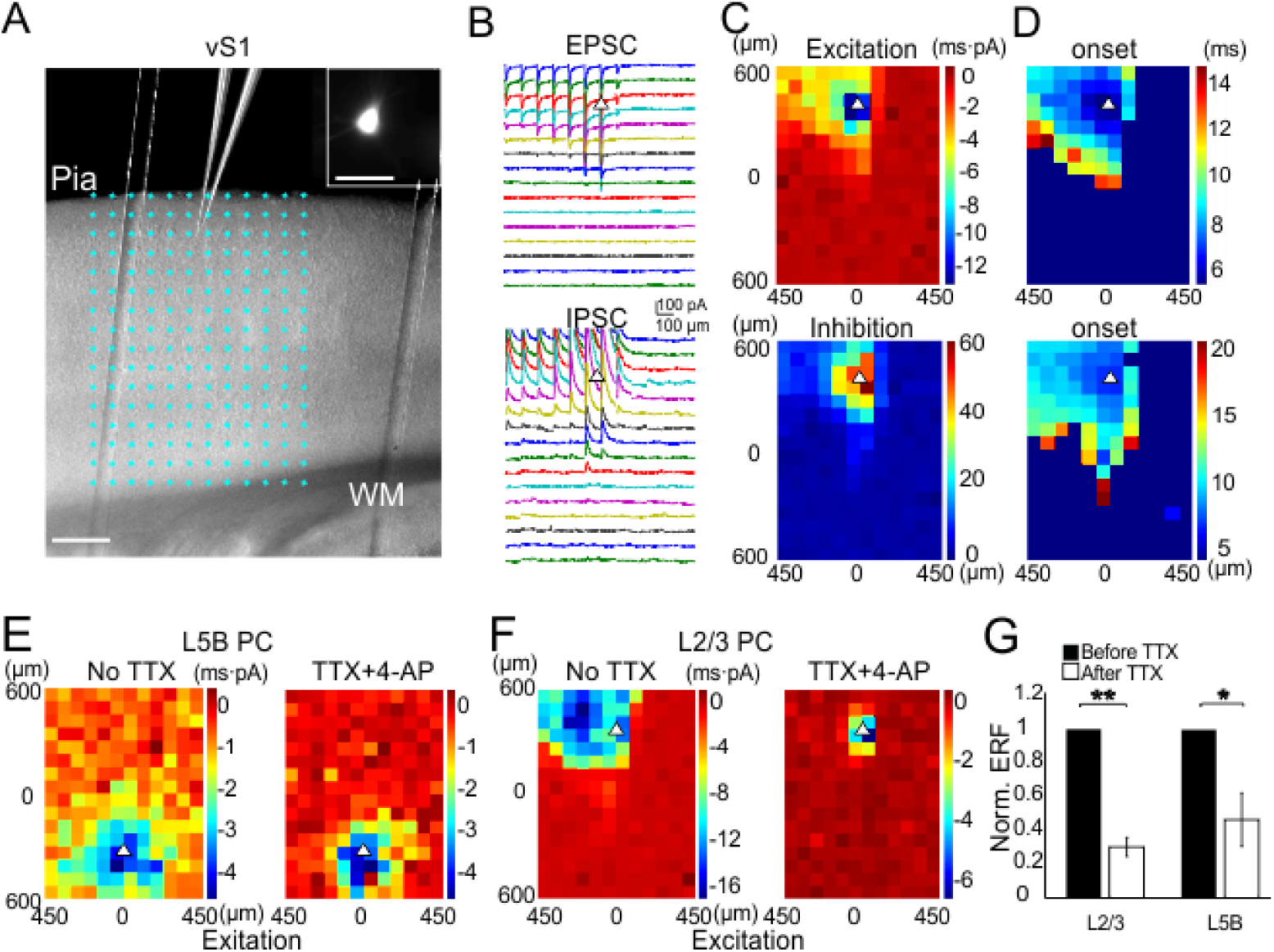
ChR2-assisted circuit mapping (CRACM) **(A)** 16 by 12 phostimulation grid overlaid with a vS1 slice showing an exemplar recording from L2/3 PC. Inset shows Alexa-594 filled neuron. (Scale bar, 200 µm). **(B)** EPSCs (top) and IPSCs (bottom) evoked by optogenetic scanning at each site; triangle indicates the soma position of recorded PC. **(C)** Charge of EPSCs or IPSCs at each site were converted to heat maps. **(D)** Heat maps show the color-coded EPSP or IPSC latency upon light stimulation at each site. **(E)** Heat maps of one L5B PC before and after TTX and 4-AP application, corresponding to polysynaptic and monosynaptic input, respectively. **(F)** Similar to e but for L2/3 PC. **(G)** Normalized REF significantly dropped after TTX and 4-AP for L2/3 PCs (P<0.01) and L5B PCs (P<0.05).

**Figure S2.**
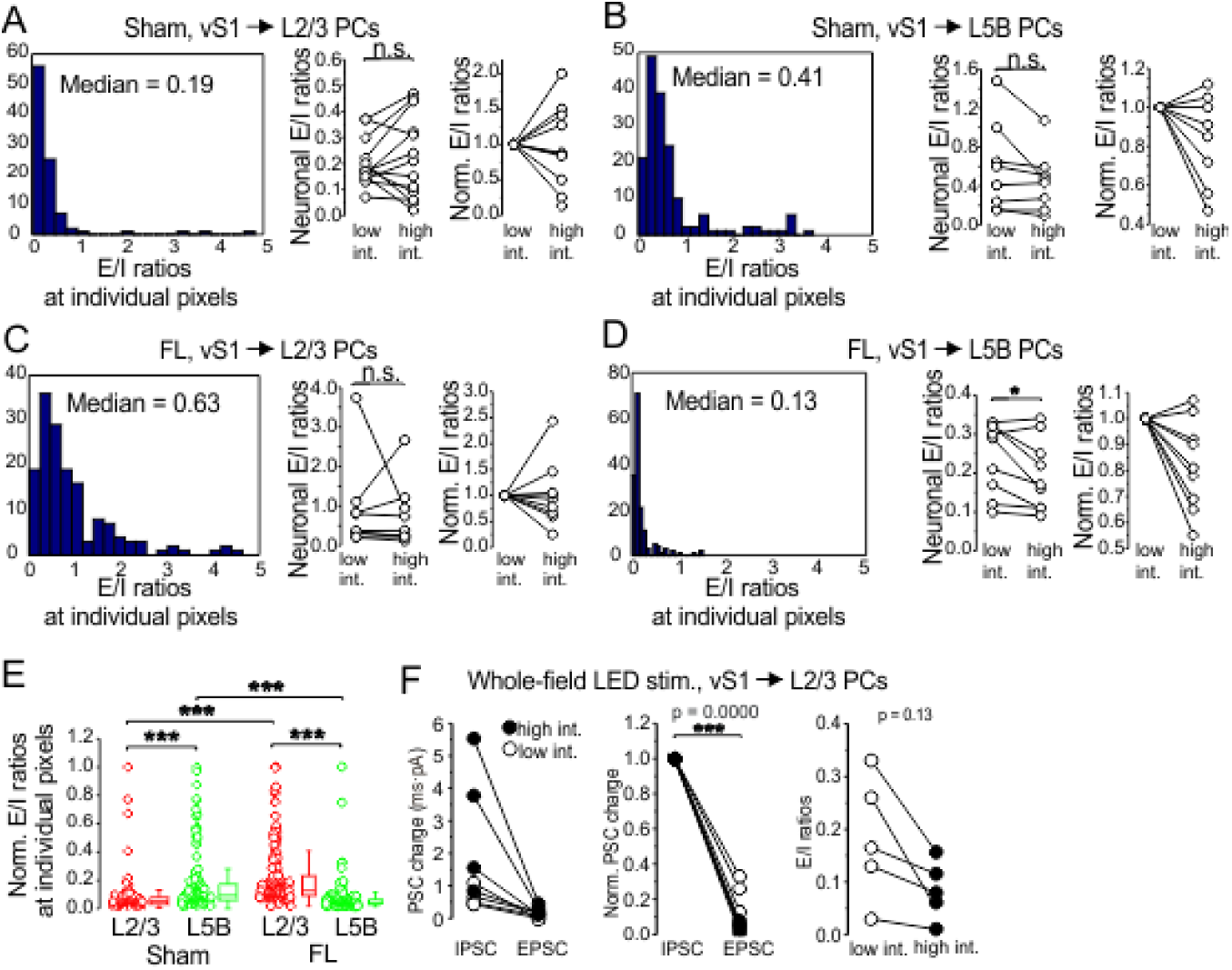
Pixel E/I ratios for vS1 → L2/3 and L5B PCs for both sham and FL groups. **(A)** Left, histogram showing the distribution of E/I ratios at individual grid pixels from 13 neurons in the sham-L2/3 group. Middle, E/I ratios under two different laser intensities (paired *t* test, *P* = 0.502, *n* = 13 neurons). Right, E/I ratios normalized to the lower intensity. **(B)** Left, histogram (n=10 neurons) for the sham-L5B group. **(C)** Left, histogram for the FL-L2/3 group. Middle: E/I ratios under two different laser intensities (paired *t* test, *P* = 0.746, *n* = 9 neurons). Right, E/I ratios normalized to the lower intensity. **(D)** Left, histogram for FL-L5B group *(n* = 13 neurons). Middle: E/I ratios under two different laser intensities (paired *t* test, *P* = 0.031, *n* = 9 neurons,). Right, E/I ratios normalized to the lower intensity. **(E)** Normalized pixel E/I ratio in L2/3 PCs in FL mice displayed significantly higher values than their counterparts in sham (red circles); meanwhile the opposite relationship was observed for L5B PCs. The pixel E/I ratios in L5B PCs were typically much higher than those in L2/3 in sham mice. However, this relationship was also reversed in FL mice. **(F)** EPSC charge was usually much higher than IPSC’s. E/I ratios were typically remained similar despite of change in stimulus strength in sham mice.

**Figure S3.**
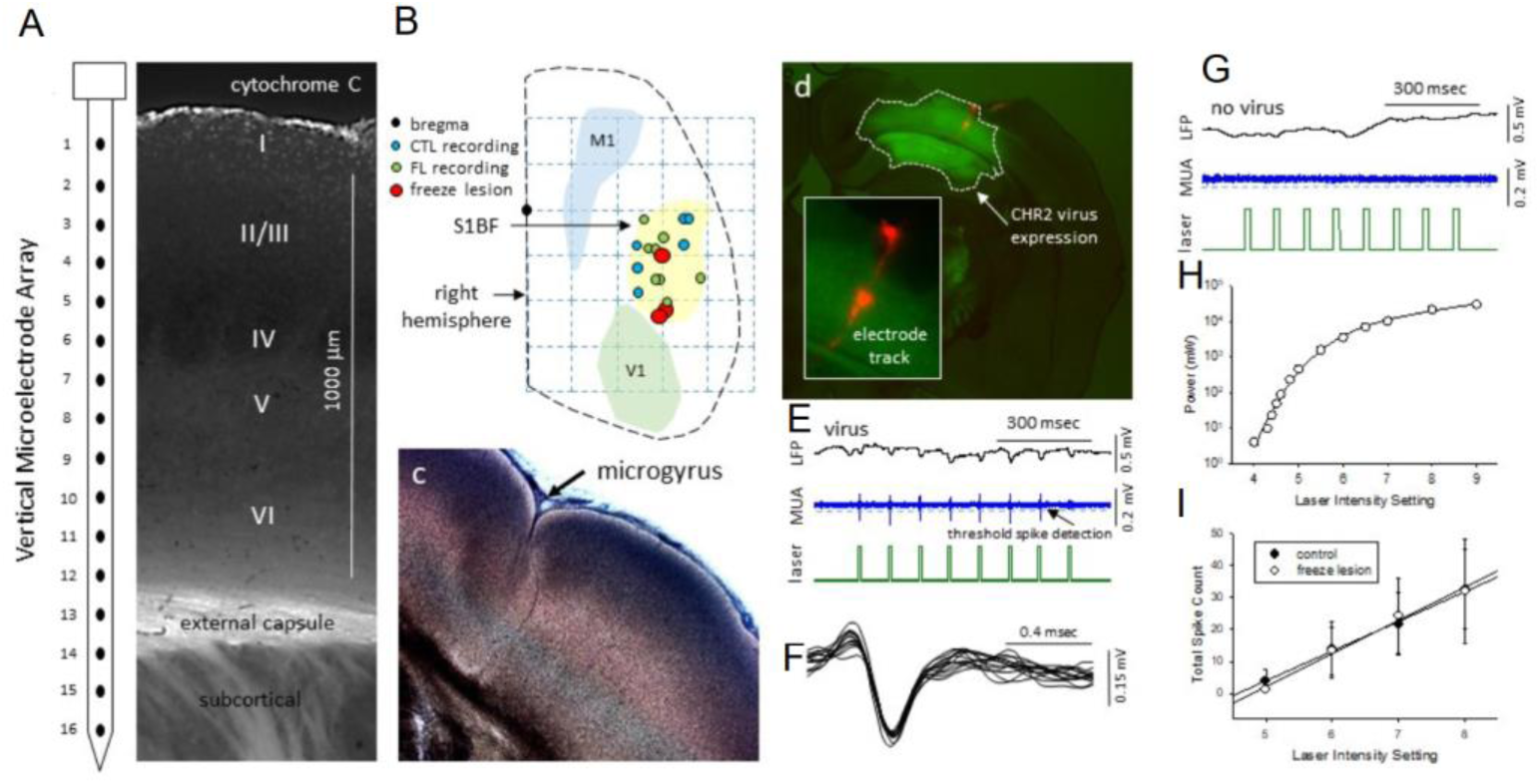
Optogenetic stimulation evoked LFP and MUA in S1BF.

**(A)** Diagram of the 16-electrode array aligned with cortical lamina of S1BF (layers I-VI) and underlying subcortical brain structures in a sham-treated mouse.

**(B)** Composite diagram indicates the location of the electrode array recording sites and FL microgyria in S1BF across all FL animals studied.

**(C)** Example of a FL microgyrus in S1BF.

**(D)** Example of AAV-CHR2-virus expression (GFP) in relation to the electrode recording sites (red, expanded in inset) in S1BF.

**(E)** Laser-evoked MUA in a mouse injected with ChR2 virus (S1BF, sham).

**(F)** Laser-evoked spikes (composite of 13 laser-evoked spikes from the same recording location).

**(G)** No laser-evoked spikes were observed in animals that did not receive ChR2 virus injection (S1BF, sham).

**(H)** Measured power values emitted from the fiber optic probe tip across laser intensity levels.

**(I)** Linear increase in total MUA spike count across intensity level for both sham (n=3, r2 = 0.997) and FL mice (n=3, r2 = 0.990). MU spike count represents average values across all 12 cortical electrodes.

**Figure S4.**
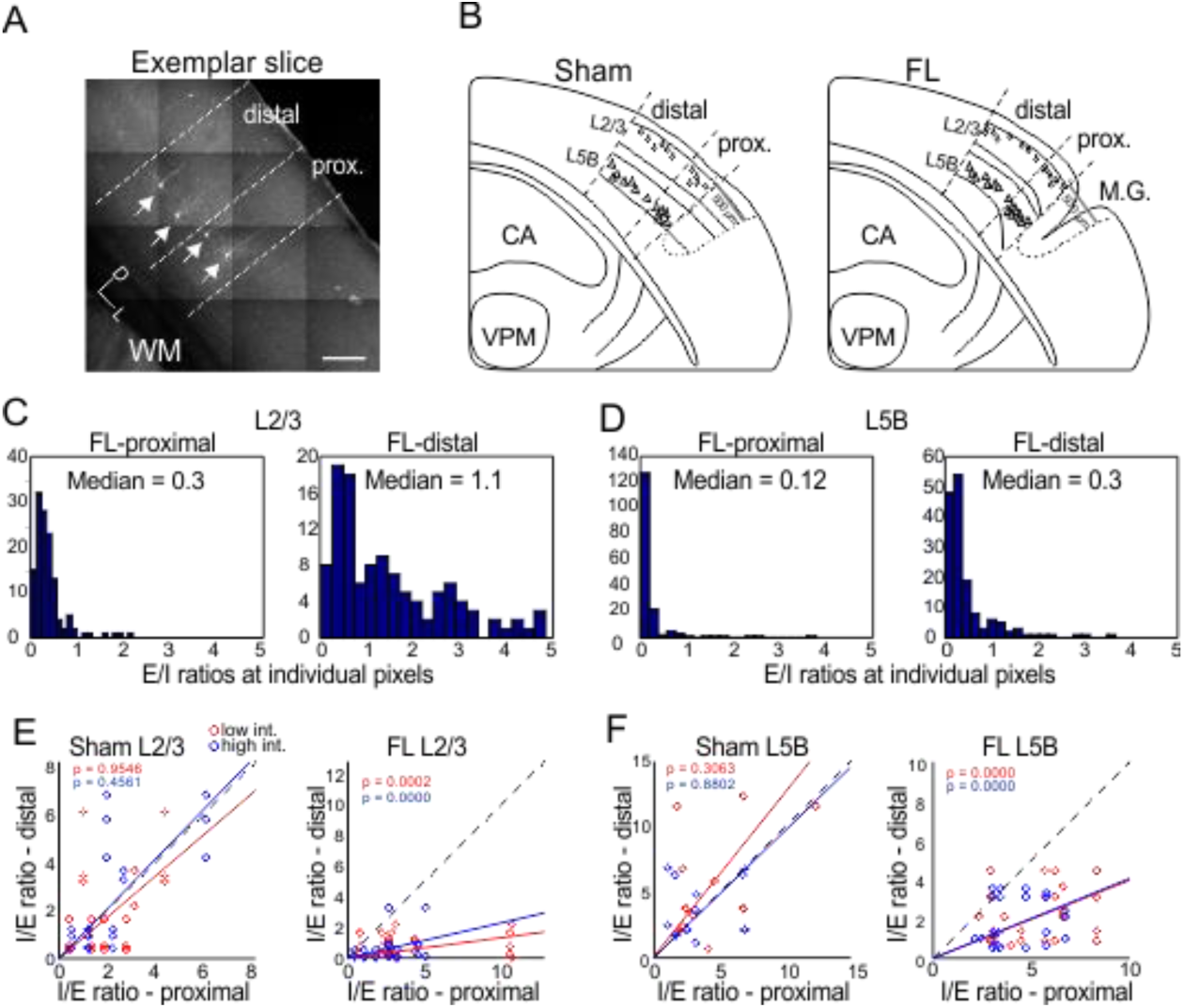
Neurons in proximal and distal FL regions differed in E/I balance. **(A)** An example slice shows four biotio-filled L5B PCs (arrows). Whether a neuron is located in proximal or distal regions depends on its relative distance away from the microgyrus. In sham mice, the center of the microgyrus was aligned with the average positions in FL mice. **(B)** Left, distribution of recorded PCs in sham mice for both L2/3 and L5B. Right, distribution of recorded PCs in FL mice for both L2/3 and L5B. **(C)** Histogram shows, in L2/3, the distribution of pixel E/I ratios for FL-proximal vs FL-distal groups. **(D)** Histogram shows, in L5B, the distribution of pixel E/I ratios for FL-proximal vs FL-distal groups. **(E)** I/E ratios (for readability) in FL-L2/3 PCs increased significantly for FL-proximal group, even under two different intensities. In sham-L2/3 PCs, this site difference was not observed. **(F)** I/E ratios in FL-proximal group were higher than that in FL-sham group. No difference was found for the sham group, as expected.

**Figure S5.**
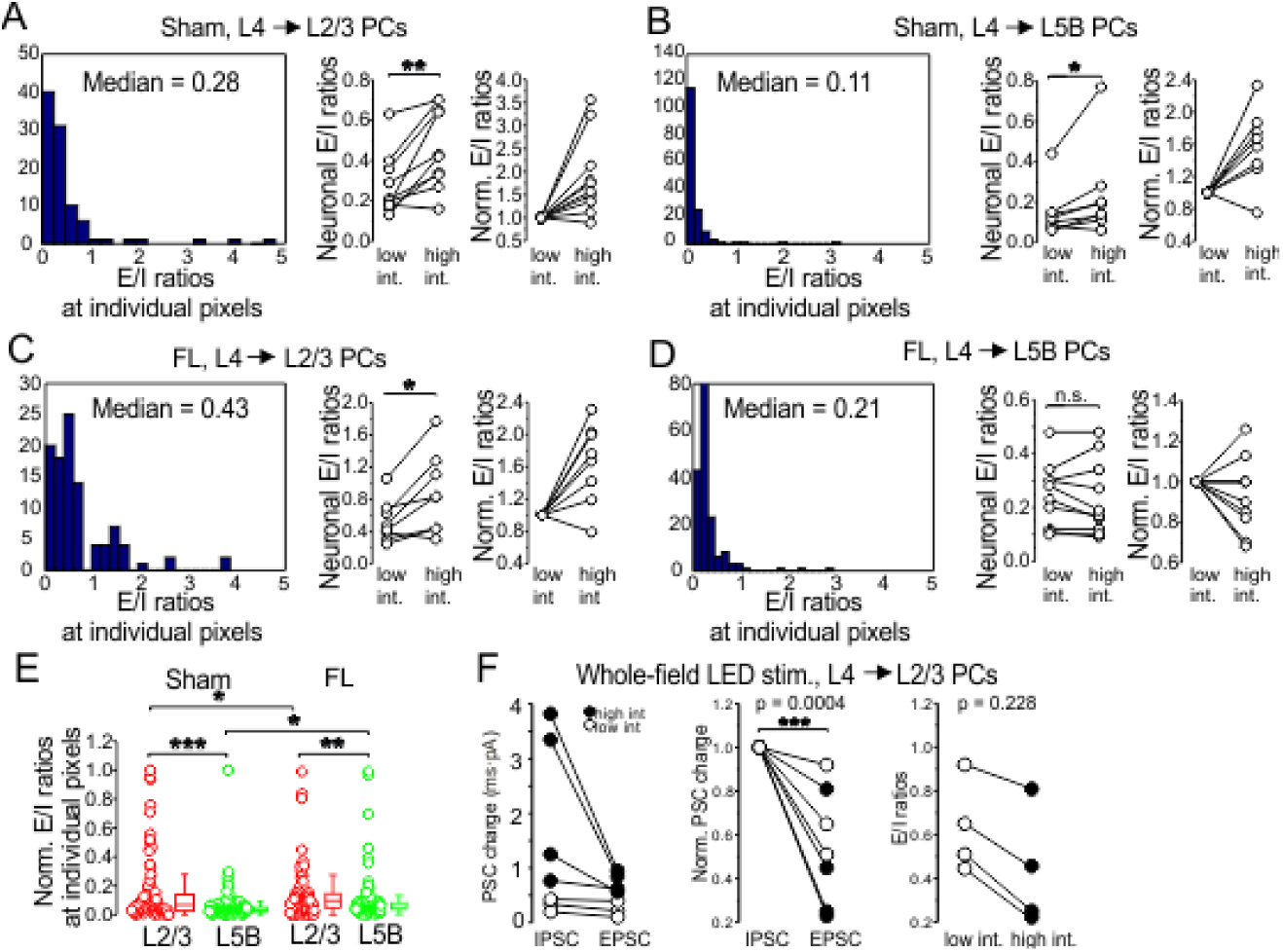
Granular layer contributes to the E/I imbalance in FL mice. **(A)** Left, histogram shows the distribution of E/I ratios at individual grid pixels from L2/3 PCs (*n* = 10, 6 mice) in sham mice. Middle: neuronal E/I ratio increased in sham-L2/3 PCs (paired *t* test, *P* = 0.002, *n* = 10 neurons, 6 mice). Right, Neuronal E/I ratios were normalized to the lower intensity. **(B)** Left, histogram showing the distribution of pixel E/I ratios of L2/3 PCs in FL mice (*n* = 10, 4 mice). Middle: neuronal E/I ratio increased at higher intensity in L5B PCs in sham mice (paired *t* test, *P* = 0.0136, *n* = 8, 4 mice). Right, Neuronal E/I ratios normalized to the lower intensity. **(C)** Histogram shows the distribution of pixel E/I ratios from L5B PCs in FL mice (*n* = 11, 4 mice). Neuronal E/I ratios increased at higher intensity (paired *t* test, *P* = 0.0472, *n* = 8 PCs, 4 mice). Right, Neuronal E/I ratios normalized to the lower intensity. **(D)** Histogram shows the distribution of pixel E/I ratios from L5B PCs in FL mice (*n* = 11, 3 mice). Neuronal E/I ratios did not differ at two laser intensities (paired *t* test, *P* = 0.5152, *n* = 8 neurons, 3 mice). Right, Neuronal E/I ratios normalized to the lower intensity. **(E)** Scatter plot shows the normalized E/I ratios at each individual pixel in all recording groups. **(F)** Left: in L2/3 PCs, EPSC charge was higher than IPSC’s. Middle: normalized EPSC charge was significantly higher than IPSC’s (P<0.001). Right, E/I ratios for L2/3 PCs remained similar with different intensities.

**Figure S6.**
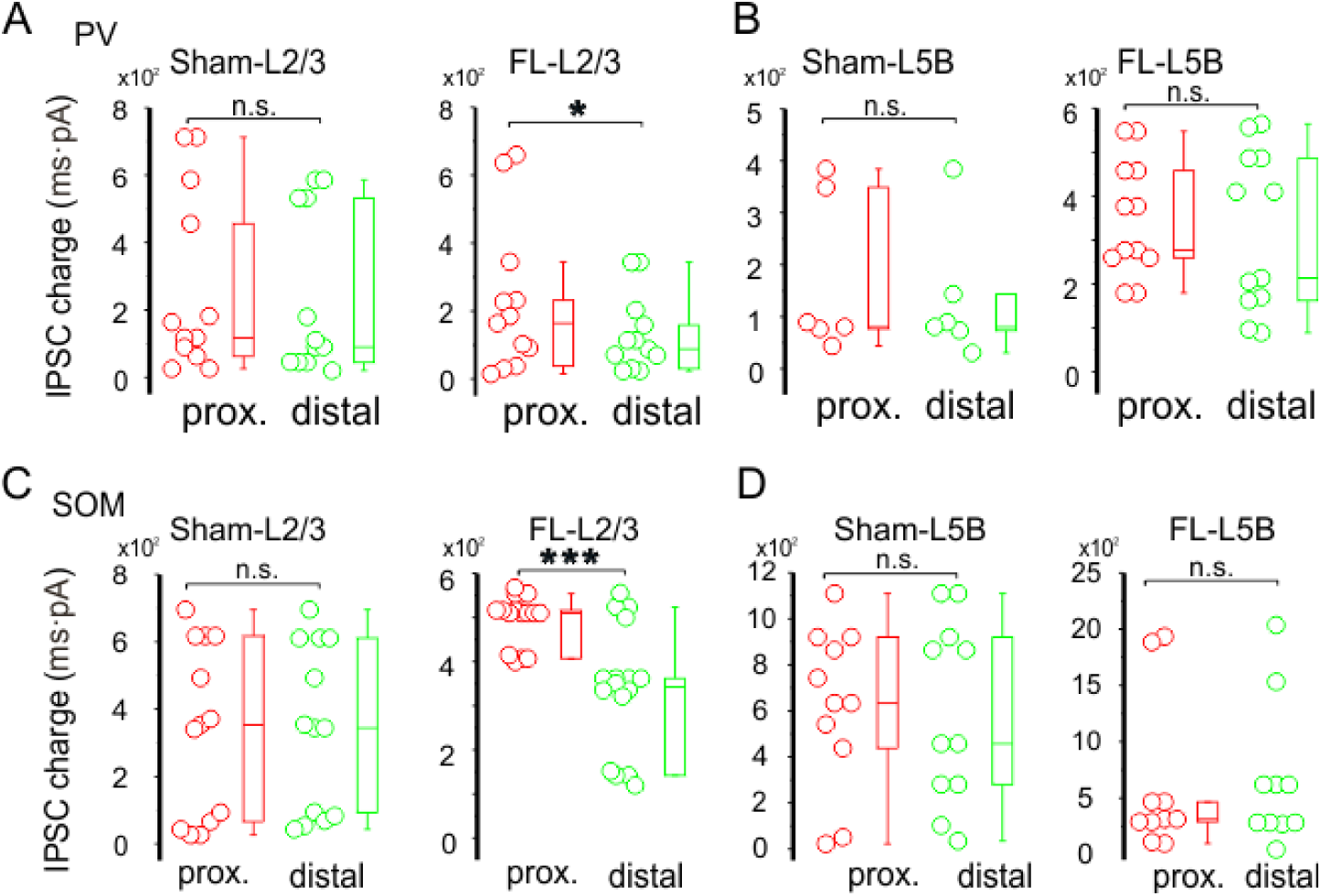
PV-vs. SOM-mediated inhibition in proximal vs. distal region. **(A)** Left, in sham L2/3 PCs, no difference between proximal and distal groups was found in IPSC charge from PV INs. Right, in FL mice L2/3 PCs, IPSC charge from PV INs was reduced in distal group (P<0.05). **(B)** In both the sham (left) and FL (right) L5B PCs, no difference between proximal and distal groups was found in IPSC charge from PV INs. **(C)** Left, in sham L2/3 PCs, no difference between proximal and distal groups was found in IPSC charge from SOM INs. Right, in FL mice L2/3 PCs, IPSC charge from SOM INs was reduced in distal group (P<0.001). **(D)** In both sham (left) and FL (right) L5B PCs, no difference between proximal and distal groups was found in IPSC charge from SOM INs.

## Notes

### Competing Interest Statement

The authors have declared no competing interest.

